# Retrieval context determines whether event boundaries impair or enhance temporal order memory

**DOI:** 10.1101/2022.01.02.474709

**Authors:** Tanya Wen, Tobias Egner

## Abstract

Meaningful changes in context create “event boundaries”, segmenting continuous experience into distinct episodes in memory. A foundational finding in this literature is that event boundaries impair memory for the temporal order of stimuli spanning a boundary compared to equally spaced stimuli within an event. This seems surprising in light of intuitions about memory in everyday life, where the order of within-event experiences (did I have coffee before the first bite of bagel?) often seems more difficult to recall than the order of events per se (did I have breakfast or do the dishes first?). Here, we aimed to resolve this discrepancy by manipulating whether stimuli carried information about their encoding context during retrieval, as they often do in everyday life (e.g., bagel-breakfast). In Experiments 1 and 2, we show that stimuli inherently associated with a unique encoding context produce a “flipped” order memory effect, whereby temporal memory was superior for cross-boundary than within-event item pairs. In Experiments 3 and 4, we added context information at retrieval to a standard laboratory event memory protocol where stimuli were encoded in the presence of arbitrary context cues (colored frames). We found that whether temporal order memory for cross-boundary stimuli was enhanced or impaired relative to within-event items depended on whether the context was present or absent during the memory test. Taken together, we demonstrate that the effect of event boundaries on temporal memory is malleable, and determined by the availability of context information at retrieval.

## Introduction

Our daily lives unfold as a continuous stream of experiences, but when we reflect on the past, we remember those experiences as distinct yet ordered events. An event is typically conceived by an observer as a segment of time at a given location that has a beginning and an end (Zacks & Tversky, 2001). A typical morning may be remembered as a series of discrete activities, such as eating breakfast, cleaning the dishes, and heading to work. Experiences are segmented into discrete events by meaningful contextual changes in one’s environment or task goals (DuBrow & Davachi, 2013; Heusser, Ezzyat, Shiff, & Davachi, 2018; Zacks, Speer, Swallow, Braver, & Reynolds, 2007). These shifts in context are known as “event boundaries”.

To study how event boundaries affect memory, researchers have devised a variety of experimental manipulations, including interleaving different stimulus classes (e.g., Clewett and Davachi, 2021; Dubrow and Davachi, 2013, 2016; DuBrow and Davachi, 2014; Sols et al., 2017), changes in perceptual context (e.g., Heusser et al., 2018; Gurguryan et al., 2021), timing context (van de Ven, Jäckels, & De Weerd, 2021), or task sets (Wang & Egner, 2022), moving through different rooms (e.g., Horner et al., 2016), and eliciting reward prediction errors (Rouhani, Norman, Niv, & Bornstein, 2020). In all of these studies, a consistent finding is that participants are better at remembering the temporal order of two items that occurred within the same event, compared to two items that appeared on either side of an event boundary. In other words, individuals are more likely to forget the temporal order of item pairs if they spanned an intervening context shift (see Clewett et al., 2019 for a review). One prominent theory for explaining this phenomenon is that sequentially presented items become linked in memory and temporal order judgments rely on reconstructing the chain of events involving the queried items (Friedman, 1993; Lewandowsky & Murdock, 1989). This account predicts that event boundaries induced by contextual changes should disrupt temporal order memory by creating a break in the chaining of events across a boundary (DuBrow & Davachi, 2013).

However, this canonical finding in the event memory literature seems to contradict many real-life scenarios, where remembering the order of experiences within an event seems more difficult than remembering the order of events. For example, it would appear much easier to recall that one had a bagel during breakfast before washing the plate during cleanup (cross-boundary) than to recall whether one took a bite out of the bagel or drank coffee first during breakfast (within-event). Additionally, people do not seem to have any trouble recalling the general trajectory of an experience that consists of multiple events and event transitions, such as a narrative of a movie (Baldassano et al., 2017; Heusser, Fitzpatrick, & Manning, 2021; Zacks, Tversky, & Iyer, 2001; Zadbood, Chen, Leong, Norman, & Hasson, 2017). In fact, participants’ recounting of narratives suggests better order memory for essential narrative elements than that of fine-scale details (Heusser et al., 2021). Intuitively, these examples seem to involve the recollection of stimulus sequences that are bound to a higher-level schema (e.g., bagel to breakfast). Additionally, the stimulus itself signals information about the event context (i.e., soapy plates are not typically seen during breakfast, but are often present during cleanup). By contrast, in typical event segmentation studies, there is no information provided about the context in which an item was encoded during memory retrieval. It is, therefore, possible that temporal order memory may, in some cases, benefit from contextual information or schematic structure afforded by event boundaries.

In line with this conjecture, some models of temporal memory would seem to predict better memory for cross-boundary item pairs. For example, “distance theories” of temporal order memory posit that recency discrimination can be based on a comparison between the strength of individual items in memory, which should be more distinct, and thus easier to discriminate, the further apart in time two items are encountered (Friedman, 1993, 2004; Hintzman, 2002). Accordingly, many studies have shown that temporal order memory improves with greater objective temporal distance between the tested items (e.g., Fortin, Agster, & Eichenbaum, 2002; Fuhrman & Wyer, 1988; St. Jacques, Rubin, LaBar, & Cabeza, 2008). Studies on event boundaries have consistently shown that items presented on either side of a contextual shift are perceived to be further apart in time (e.g., Bangert, Kurby, Hughes, & Carrasco, 2019; Ezzyat & Davachi, 2014; Lositsky et al., 2016). Following this line of reasoning, one would predict that an event boundary should facilitate temporal order judgments, as items crossing a boundary would be perceived as having a greater distance between them. This was recently demonstrated in a model simulation (Rouhani et al., 2020), where better temporal order memory was predicted for items spanning an event boundary. However, this prediction runs counter to empirical findings from the event segmentation literature.

In the current study, we conducted a series of four behavioral experiments to reconcile these disparate intuitions concerning the effects of event boundaries on temporal order memory. We had participants encode series of stimuli associated with periodically changing contexts and tested their temporal order memory for pairs of stimuli. Based on the argument developed above, we hypothesized that it should be possible to obtain superior order memory for cross-boundary items if those items provide information about the encoding context at retrieval (as in the eating/cleaning example). To this end, we developed two novel event memory paradigms that created conditions where either all (Experiment 1) or a subset of (Experiment 2) the task stimuli inherently signaled their encoding context, by making those stimuli uniquely applicable to their respective encoding task. In line with our predictions, when stimuli provided information about their encoding context at retrieval, we observed a “flipped” order memory effect, whereby temporal memory was superior for cross-boundary than within-event item pairs. In Experiments 3 and 4, we added context information at retrieval to a more standard event memory protocol where stimuli are not inherently associated with a specific encoding context but are combined with arbitrary context cues (colored frames, see Heusser et al. 2018). We found that whether temporal order memory for cross-boundary stimuli is enhanced or impaired relative to within-event items did indeed depend on whether the context was present or absent during the memory test. Taken together, we present compelling evidence that the nature of the effects that event boundaries have on temporal order memory is determined by the availability of event context information at retrieval.

## General Methods

### Participants

Participants were recruited through the Duke Department of Psychology and Neuroscience Subject Pool for course credit, and through Amazon Mechanical Turk (MTurk) for monetary compensation. All experiments were conducted online. For each experiment or experimental group, we recruited ~30 participants (this number was doubled in Experiment 3 due to having half the number of runs in each condition). The sample size was chosen based on prior studies on event boundary-related memory effects (power = 0.8, α = 0.05 for 26 subjects; Dubrow & Davachi, 2013; Heusser et al., 2018). The experiment was approved by the Duke University Institutional Review Board. Informed consent was obtained from all participants prior to their participation.

### Procedure

In each experiment, participants completed 10 task runs preceded by 1 practice run. Each run consisted of (1) an encoding phase, (2) a delay phase, and (3) a memory phase.

On each trial of the encoding phase, participants were asked to make a decision about a target stimulus shown in the center of the screen (the precise tasks differed in each experiment and will be described in more detail in each of their corresponding sections). There were two response choices displayed in the bottom left and bottom right, and participants used the “Z” and “M” keys to select the left or right answer, respectively. Each trial of the encoding phase consisted of a 3.5 s target display followed by a 1 s fixation. Participants were encouraged to be as accurate as possible. We did not provide trial-by-trial feedback, as we did not want occasional negative feedback to potentially create event boundaries, but we provided participants with feedback on their proportion of correct responses after each run. Each encoding phase included several changes in context. Context was operationalized as a common task feature that remained the same throughout several consecutive trials of each run (Howard & Kahana, 2002; Manning et al., 2013). Event boundaries were created by an abrupt change in the shared task feature (e.g., Heusser et al., 2018; Wang & Egner, 2022).

Each encoding phase was followed by a distractor task to create a brief delay between the memory encoding and retrieval phases. Specifically, in the delay phase, participants were given an alphanumeric string between 4~6 characters long (e.g., “Wi54b”) and were asked to type it backwards (e.g., “b45iW”) into a text box. They were able to proceed after correctly completing the task or were asked to try again.

After the delay phase, participants completed the memory phase where they were tested for their temporal memory for images from the preceding encoding phase. On each trial of the temporal memory tests, participants were presented with a side-by-side pair of two images, and asked about their order (“Please select the image that was seen first”) and perceived temporal distance (“How far apart in time were these images presented?”). Temporal distance was rated on a scale of “very close,” “close,” “far,” and “very far,” and the ratings were converted to a numerical scale (1-4) for analysis. Temporal memory tests included image pairs that shared the same context (within-event) or were from neighboring contexts and thus spanned an event boundary (cross-boundary). Within each experiment, all image pairs regardless of condition had the same number of intervening items during encoding, and therefore identical objective temporal distance. In Experiment 4, participants additionally completed source memory tests in each run, where they were asked to select the context that was previously presented with each queried stimulus. All memory trials were self-paced.

Before the beginning of each experiment, there was a practice run to ensure that participants understood the task. Participants were required to perform above 80% accuracy in the encoding task and above 50% accuracy in the temporal order memory task (and additionally, in Experiment 4, above 50% accuracy in the source memory task) in the practice run before proceeding to the main experiment. Otherwise, they were to repeat the practice.

### Data analysis

For the encoding phase data, we examined accuracy and reaction time. We excluded reaction times for outliers greater than 3 standard deviations from the mean of each phase or memory test, as well as for trials associated with an incorrect response. For memory phase data, we analyzed accuracy rates for temporal order judgments and distance ratings from subjective temporal distance estimates using t-tests and/or ANOVAs, depending on the experiment. While our paper focuses on memory accuracy and distance ratings, as these two measures have been considered charateristic measures of event boundary effects (Clewett, DuBrow, & Davachi, 2019), we report details of reaction times during the memory tests in the Supplementary Material. Correction for multiple comparisons was performed using the false discovery rate (FDR; Benjamini & Hochberg, 1995) for each analysis.

### Data and code sharing

All data, experimental stimuli, and task/analysis code are available at https://github.com/tanya-wen/Event-Boundary.

## Experiment 1

In Experiment 1, we sought to experimentally capture the type of real-life scenario alluded to in the Introduction, where stimuli are strongly associated with specific contexts. To this end, we exposed participants to discrete and unrepeated tasks, where each trial in the event shared an overarching theme. This included tasks like categorizing images, filling in words, solving math problems, solving puzzles, judging the veracity of statements, etc. (Fig. 1A). Thus, every event boundary involved a shift both in the task and the stimulus material. This served to accentuate the uniqueness of each event and maximize contextual differences between the stimuli, as prior work has shown that changes to both the stimulus class and the task influence the organization of memory (Polyn, Natu, Cohen, & Norman, 2005). Because the stimuli presented in a given task were unique to that task, each stimulus inherently provided information about its encoding context during the memory test. We tested temporal order memory via recency discrimination for pairs of items presented within the same event and across boundaries. On one hand, it is possible that event boundaries would interrupt the chaining across stimuli, and we would observe the classic effect of within-event item pairs showing better temporal order memory than cross-boundary pairs (DuBrow & Davachi, 2013). However, we hypothesized that with the current design, where the stimuli provide event context information during memory retrieval, cross-boundary pairs would be associated with better temporal order memory than within-event pairs.

**Figure 1.**
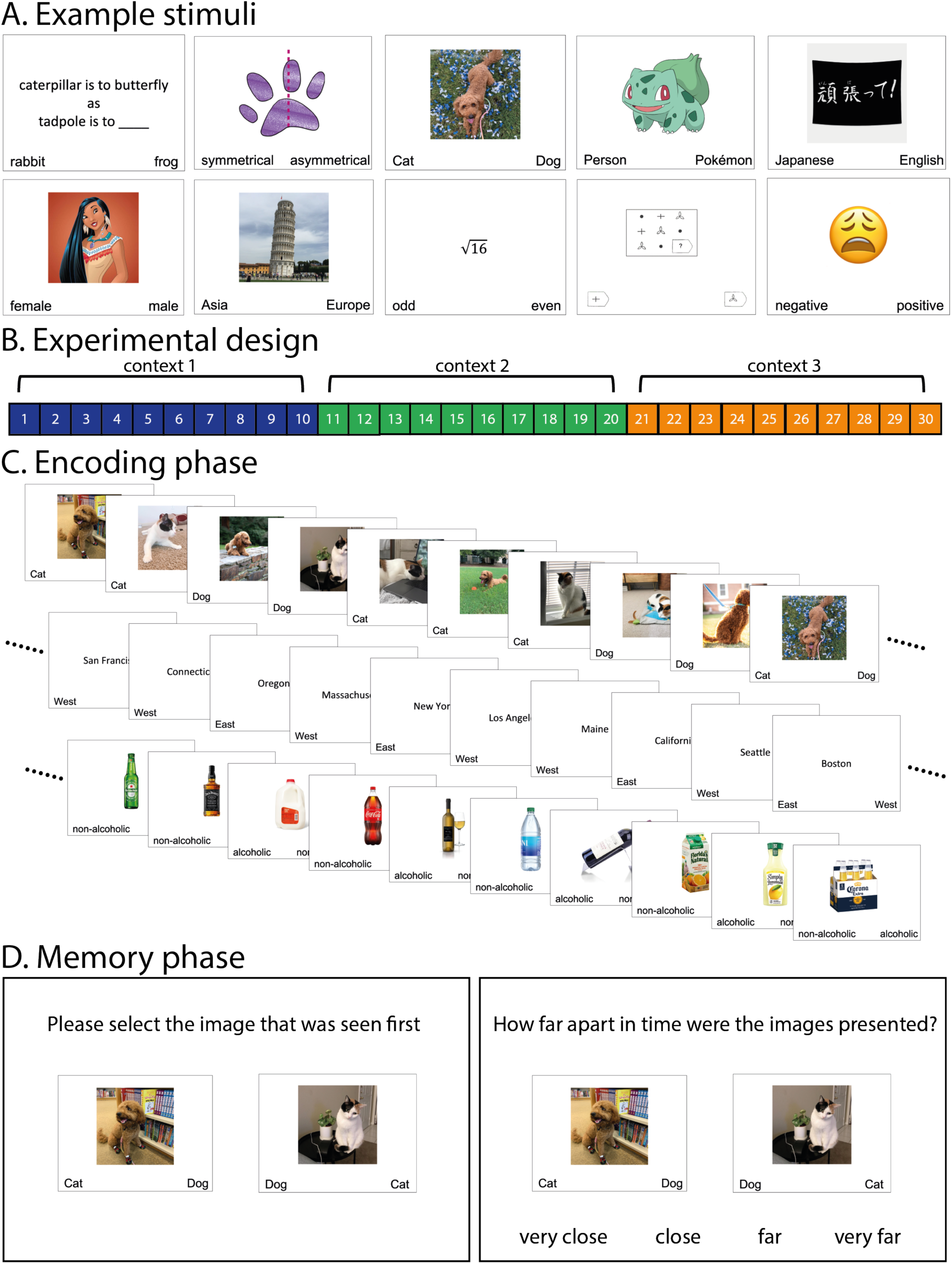
(A) Example stimuli used in Experiment 1. (B) Experimental design. Each run consisted of three events where participants had to perform ten trials of a particular task. (C) Example encoding phase. (D) Memory phase. Each trial consisted of a temporal order memory discrimination of two stimuli, followed by a judgment of their temporal distance.

### Methods

#### Participants

31 participants (6 male, 23 female, and 2 did not wish to reply; ages 18~22, mean = 19, SD = 1.13) were recruited from the Duke SONA subject pool for 1.5 course credits. 1 additional participant was excluded due to performance accuracy below 80% during the encoding task.

#### Stimuli

33 different task contexts were generated by the experimenter (see Figure 1A for examples). Each context was only encountered once during the experiment, and included a diverse range of two-alternative choice tasks. For example, participants may be asked to complete analogies, judge whether a shape is symmetrical, categorize images and words, solve math problems, complete matrix reasoning tasks, judge emotions, etc.

#### Procedure

Figure 1B shows an illustration of the experimental design and Figure 1C shows an example of the encoding phase. During this phase, each run consisted of 30 trials, divided into 3 task contexts, with each context consisting of 10 trials where participants performed a particular two-alternative choice task. Context and stimulus order were randomized across subjects, and there were an equal number of left and right correct answers.

Figure 1D illustrates the memory phase. Each trial consisted of a temporal order memory discrimination of two stimuli, followed by a judgment of their temporal distance. To obtain memory effects as a function of serial position during encoding, we tested all possible pairs of stimuli that were spaced with 2 interleaving trials during encoding. There were 27 memory trials in total per each run, and they were presented in random order to the participants.

### Results

#### Encoding

Figures 2A and 2B show the encoding task accuracy and response times for each item position, averaged across events. Accuracy across all trials was high (mean = 93.24%, SD = 5.29%). There were some differences in pairwise comparisons of accuracy based on encoding position, mainly with the first trial of a new task context being lower in accuracy. Additionally, the first trial of a new task context had the longest reaction time compared to the remaining trials (all ts > 7.27; all ps < 0.001). There was no difference in reaction times among trials at positions 2~10 (all |t|s < 2.10; all ps > 0.18). These results reflect the classic task switch cost (e.g., Rogers & Monsell, 1995), whereby people are slower and more error-prone when switching to a new task compared to repeating a task set.

**Figure 2.**
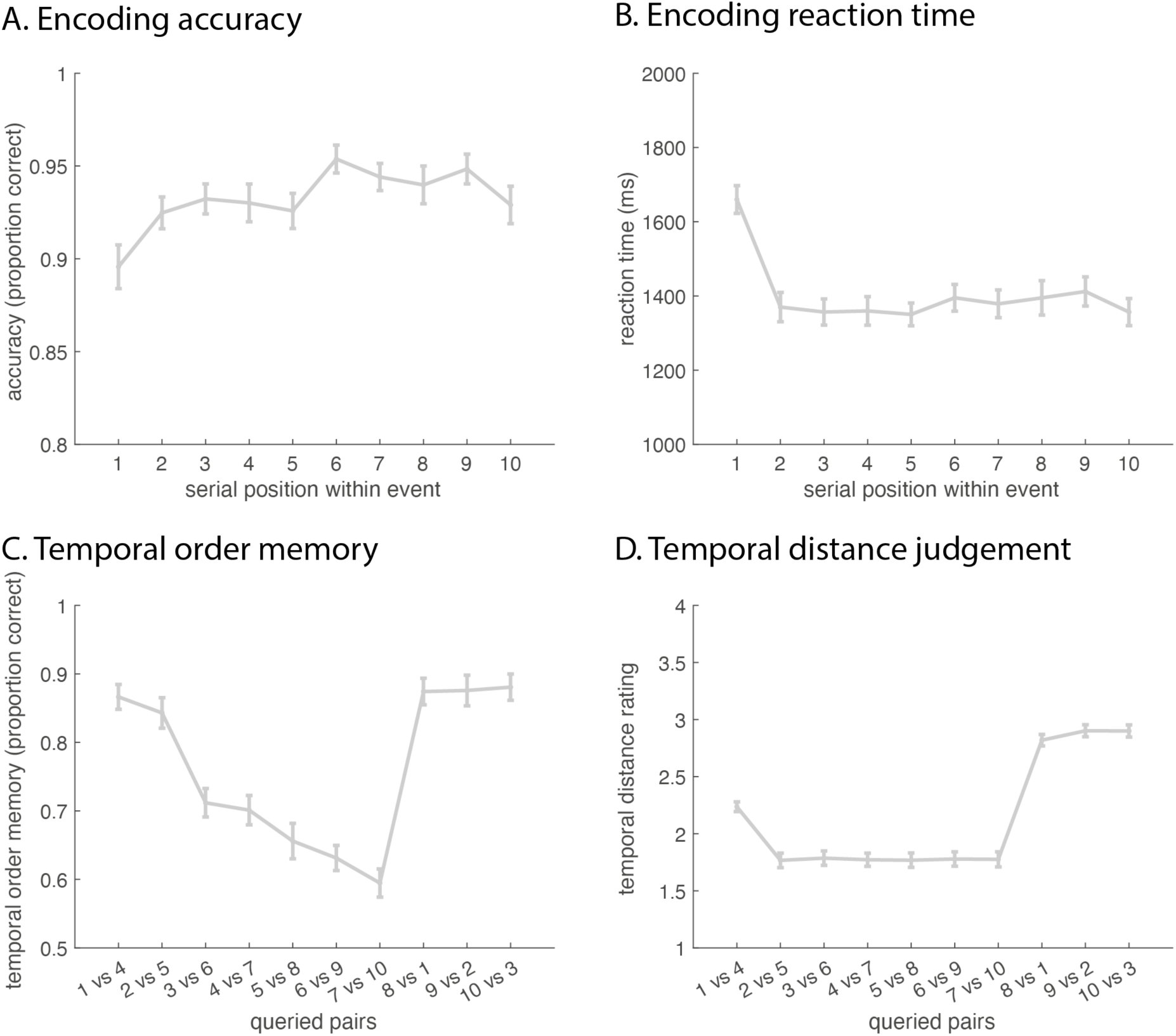
Experiment 1 encoding task accuracy (A) and response times (B) averaged across participants and plotted as a function of trial position within an event. (C) Temporal order memory and (D) temporal distance judgment as a function of encoding serial position. Error bars represent standard error.

#### Temporal memory

Figure 2C illustrates temporal order memory as a function of serial order during encoding. We found that accuracy as a function of serial position followed a U-shaped curve. Pairwise comparisons showed significant differences between numerous queried pairs (summarized in Supplementary Table 1). To point out the most prominent effects, we observed that among within-event items, order memory for early pairs, (1 vs. 4) and (2 vs. 5), was significantly more accurate than for later pairs (3 vs. 6), (4 vs. 7), (5 vs. 8), (6 vs. 9), and (7 vs. 10) (all ts > 7.30, all ps < 0.001). There was a decline in order memory for within-event pairs that appeared later in encoding, with several later pairs showing gradually worse memory. Finally, all cross-event pairs showed better temporal order memory than within-event pairs, except for the earliest two (all ts > 7.52, all ps < 0.001). Combing all within-event (mean = 71.49%, SD = 9.23%) and cross-event pairs (mean = 87.69%, SD = 10.55%), we still find that cross-event pairs had better temporal order memory (t = 12.99, p < 0.001). The mean reaction time for correct trials was 2570.17 ms (SD = 796.22 ms), and there was no significant difference in reaction time between within-event and cross-boundary pairs (t = 1.67, p = 0.11; Supplementary Figure 1A).

Figure 2D illustrates temporal distance judgment as a function of encoding serial order (summarized in Supplementary Table 2). We found the first within-event pair of a new task context (1 vs. 4) was significantly different from all other pairs (all |t|s > 10.13, all ps < 0.001), being perceived as further apart than other within-event pairs but closer together than cross-boundary pairs. There were no differences among the remaining within-event pairs (all |t|s <= 1.48, all ps > 0.22), which were all judged to be closer in time than all cross-boundary pairs (all ts < −12.70, all ps < 0.001). Combing all within-event (mean = 1.84, SD = 0.32) and cross-event pairs (mean = 2.87, SD = 0.28), we find that cross-event pairs were judged to be more distant (t = 14.12, p < 0.001). In regards to reaction time, participants were overall slower in judging cross-boundary pairs (mean = 1888.89 ms, SD = 657.22 ms) compared to within-event pairs (mean = 1617.26 ms, SD = 484.37 ms; Supplementary Figure 1B).

### Discussion

Our primary interest for Experiment 1 was to examine whether distinctive task contexts with unique stimulus sets that carry contextual information during the memory phase could cause a “flip” in the classic temporal order memory effect, resulting in better order memory for stimuli spanning two events compared to those that shared the same event. Indeed, contrary to the canonical finding in a large event memory literature (e.g., DuBrow & Davachi, 2013; DuBrow & Davachi, 2014; DuBrow & Davachi, 2016; Sols et al., 2017; Clewett & Davachi, 2021; Gurguryan et al., 2021; Heusser et al., 2018; Wang & Egner, 2022; Horner et al., 2016; Rouhani et al., 2020), our results show on average better temporal order memory for cross-event pairs. We further showed, by systematically testing each possible pair throughout the run, that order memory for early within-event pairs was better than that for later within-event pairs. Our temporal distance judgment results were consistent with prior findings, showing that item pairs crossing an event boundary are perceived to be further apart in time (Ezzyat & Davachi, 2014). We additionally found that the first item pair of a new task event was perceived to be further apart in time than other within-event pairs, yet closer together than cross-event pairs. In sum, the main findings of Experiment 1 provide initial support for the proposal that, under conditions of strong item-context associations, event segmentation results both in enhancing the perceived distance of cross-boundary events and in improved differentiation of the order in which those items were encoded. This is expected based on temporal distance models of order memory (Friedman, 1993, 2004; Hintzman, 2002) but runs counter to the canonical findings in event boundary studies. In Experiment 2, we probed whether we could produce both the traditional and flipped order memory patterns within a single task, as a function of whether cross-boundary stimuli clearly belonged to distinct contexts or not.

## Experiment 2

In Experiment 1, we showed a “flip” of the classic event boundary effect on temporal order memory under conditions where encoding tasks were unique, and both the task and the type of stimuli changed between events. This served to accentuate the distinction between successive contexts and resulted in stimuli signaling their unique context during memory retrieval. This differs from previous event segmentation studies where task contexts are typically not unique and the same categories of stimuli are encountered across different contexts. For instance, many studies involved alternating between two task contexts in an ABAB design (e.g., DuBrow & Davachi, 2013; Sols et al., 2017; Wang & Egner, 2022). Other studies, while having unique contexts each run, were similar in the overarching task and employed the same type of target stimuli throughout the experiment (e.g., Gurguryan et al., 2021; Heusser et al., 2018), such as judging whether a particular color would look pleasant on a grayscale object. Notably, it has been shown that categories can help organize memory (Bousfield, Cohen, & Whitmarsh, 1958; Polyn et al., 2005; Tzeng & Cotton, 1980). The set-up in Experiment 1 results in stimulus categories signaling their encoding context at retrieval, whereas in prior event memory studies target stimuli did not provide contextual information at retrieval. While this comparison between Experiment 1 and prior studies is suggestive, to corroborate that the flipped order memory results of Experiment 1 are attributable to this design difference, one would ideally demonstrate both the classic and the flipped effect of order memory for cross-boundary items in a single experiment where the only difference between conditions is whether event changes involve a task change only or both a task and stimulus category change.

In Experiment 2, we designed a task with hierarchical event boundaries, where lower-order transitions involved a change in task rule only (e.g., Wang & Egner, 2022; Heusser et al., 2018; Gurguryan et al., 2021; Horner et al., 2016; Clewett et al., 2021) and higher-order transitions involved a change in both the task rule and stimulus category (Experiment 1; see also Crittenden, Mitchell, & Duncan, 2015; Smith, Mitchell, & Duncan, 2018). We hypothesized that stimulus pairs crossing only task transitions would be impaired in their temporal order judgment compared to within-event pairs, consistent with prior studies. By contrast, we hypothesized that stimulus pairs crossing task and stimulus category changes would show enhanced temporal order memory, following the results of Experiment 1.

### Methods

#### Participants

A new set of 28 participants (6 male, 22 female; ages 18~21, mean = 18.93, SD = 0.77) were recruited from the Duke SONA subject pool for 1.5 course credits. 1 additional participant was excluded due to failure to engage in the task.

#### Stimuli

Stimuli consisted of 112 objects (28 small manmade, 28 small natural, 28 large manmade, 28 large natural), 112 words (28 1-syallable abstract, 28 1-syallable concrete, 2-syallable abstract, 28 2-syallable concrete), and 112 scenes (28 indoor cottage, 28 indoor skyscraper, 28 outdoor cottage, 28 outdoor skyscraper). The object and scene images were obtained using Google Image Search. The word stimuli were taken from frequent words in the MRC Psycholinguistic Database (Coltheart, 1981).

#### Procedure

In this experiment, participants were instructed to perform a total of 6 tasks. For objects, they were asked to judge either their origin (manmade or natural) or size (small or large; participants were told to use a shoebox as reference); for words, they were asked to judge either their abstractness (abstract or concrete) or the number of syllables (1 or 2); and for scenes, participants were asked to judge either their location (indoor or outdoor) or building type (cottage or skyscraper).

Figure 3 provides an illustration of the experimental paradigm. During the encoding phase, participants were asked to categorize the center stimuli for 5 consecutive trials according to one task before switching either task only (e.g., from judging object origin to judging object size), or both task and stimulus category (e.g., from judging object origin to judging word abstractness). Each run consisted of 3 higher-order and 6 lower-order contexts during encoding. Context and stimulus order were newly pseudo-randomized across participants in each run, such that each pair of lower-order contexts consisting of the same stimulus category were adjacent events.

**Figure 3.**
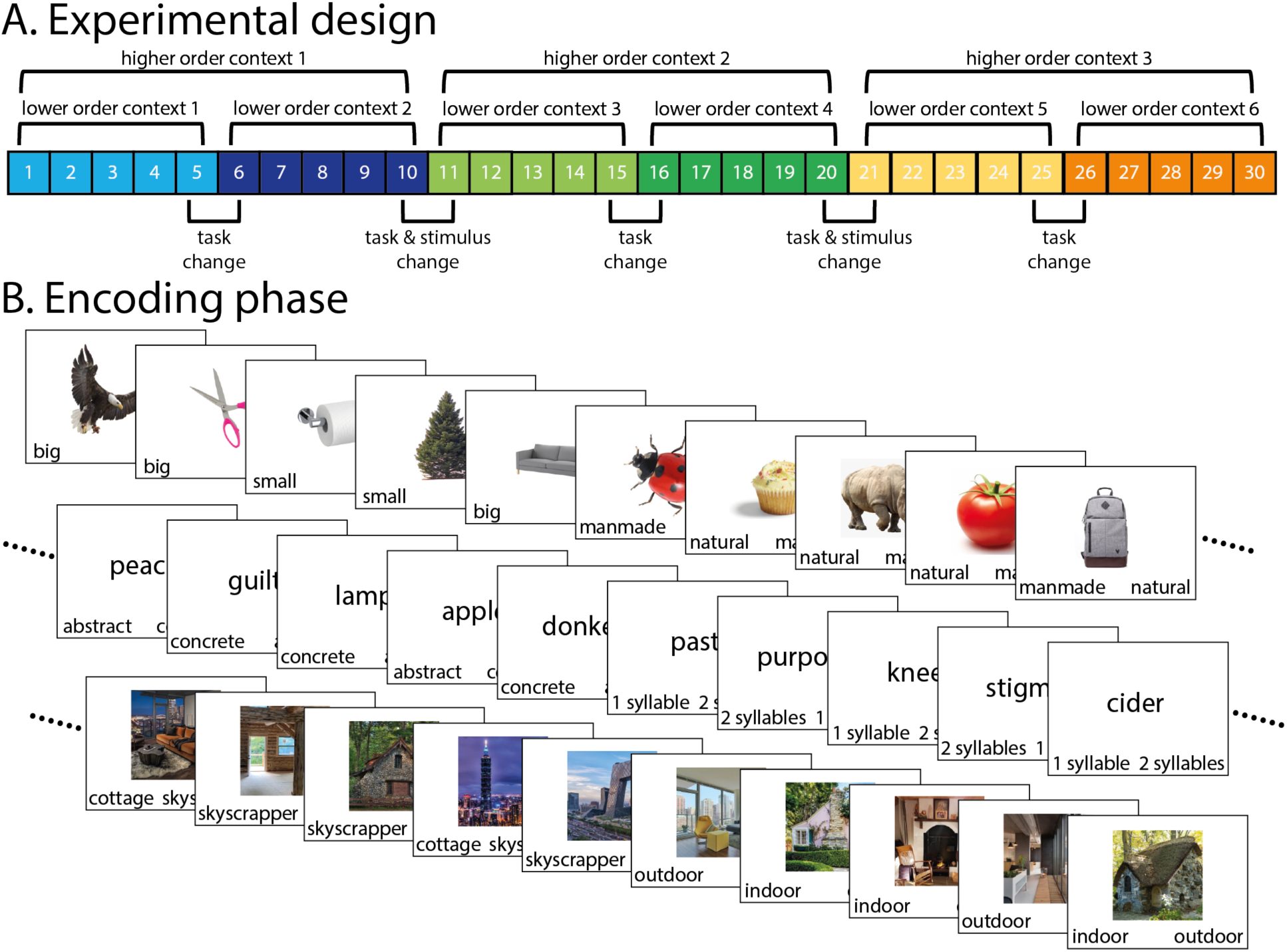
(A). Experiment 2 design, illustrating the ordering of task, and task and stimulus category changes. (B). Example encoding phase. There were three stimulus categories in each run (objects, words, and scenes), with each stimulus category consisting of two task rules.

For the temporal memory phase, we tested temporal order memory followed by subjective temporal distance judgment for each possible pair (same as in Experiment 1).

### Results

#### Encoding

Figures 4A and 4B show the mean encoding task accuracy and response times at each serial position, averaged across events. Accuracy across all trials was high (mean = 87.20%, SD = 6.87%). The 6th and 7th trials were higher in accuracy than the 2nd trial, possibly due to a speed-accuracy trade-off. The 1st and 6th trials within a higher-order event (corresponding to task switch trials) had the longest reaction time compared to the remaining trials (all ts > 2.68; all ps < 0.03), with the exception of no significant difference between the 1st and 7th trial (t = 1.56, p = 0.16). Additionally, reaction times at positions 2, 3, 4 were faster than positions 7, 8, 9, 10 (all ts > 2.27, all ps < 0.05).

**Figure 4.**
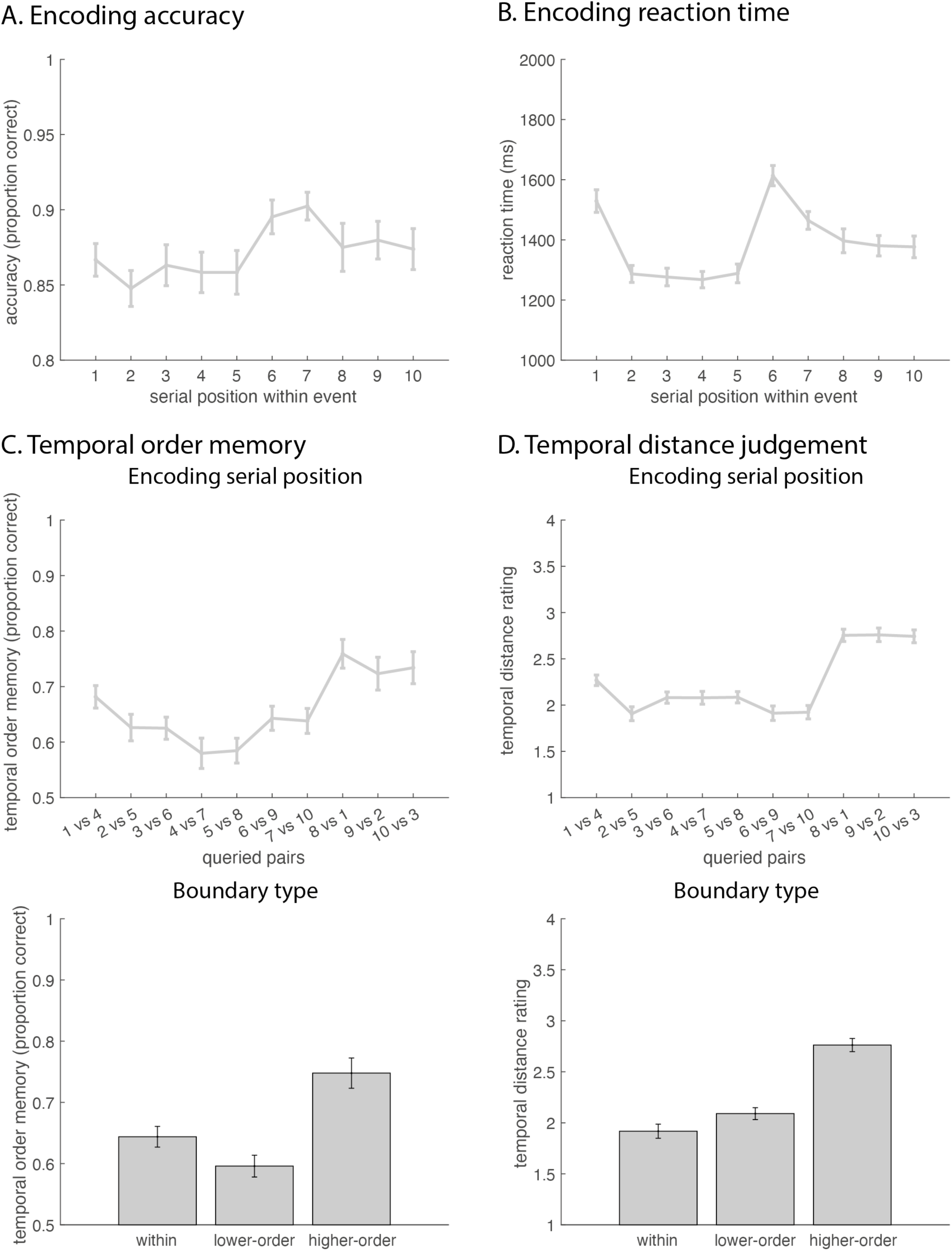
Experiment 2 encoding task accuracy (A) and response times (B) averaged across participants and plotted as a function of trial position within an event. Temporal order accuracy (C) and temporal distance judgment (D) as a function of trial position pairs (top) and boundary type (bottom; within-event, task switch, and task and context switch). Error bars represent standard error.

#### Temporal memory

Figure 4C illustrates temporal order memory accuracy as a function of serial position and boundary type (within-event, lower-order boundary, or higher-order boundary). The full range of pairwise comparisons is summarized in Supplementary Table 3. We found that accuracy and serial position follow a U-shaped curve. Temporal order of stimuli spanning a lower-order boundary (task switch only; mean = 59.60%, SD = 9.42%) was more poorly recalled than that of stimuli within the same event (mean = 64.39%, SD = 8.96%; t = −4.19, p < 0.001) as well as of stimuli spanning a higher-order boundary (task and stimulus switch; mean = 74.78%, SD = 13.10%; t = −8.50, p < 0.001). Temporal order memory for pairs within an event was also worse than for those spanning a higher-order boundary (t = −5.16, p < 0.001). In terms of reaction times (Supplementary Figure 2A), participants were faster at recalling the temporal order of stimuli spanning a higher-order boundary (mean = 2388.80 ms, SD = 1043.45 ms) compared to stimuli within the same event (mean = 2688.85 ms, SD = 1218.05 ms; t = 3.42, p < 0.01) as well as those spanning a lower-order boundary (mean = 2792.67 ms, SD = 1109.84 ms; t = 3.00, p < 0.01). There were no differences in reaction time for stimuli within the same event and those spanning a lower-order boundary (t = −0.93, p = 0.36).

Figure 4D shows temporal distance judgments as a function of serial position and boundary type. The full range of pairwise comparisons is summarized in Supplementary Table 4. We found that the first within-event pair following a boundary (1 vs. 4) was significantly different from all other pairs (all ts > 4.99, all ps < 0.001), being perceived as further apart than other within-event pairs but closer together than cross-boundary pairs. Moreover, stimuli within the same event (mean = 1.92, SD = 0.37) were perceived to be closer in time than those spanning a lower-order boundary (task switch only, mean = 2.09, SD = 0.31; t = −5.27, p < 0.001), as well as a higher-order boundary (task and stimulus switch, mean = 2.76, SD = 0.34; t = −9.92, p < 0.001). Pairs spanning a lower-order boundary were additionally perceived to be closer in time than those spanning a higher-order boundary (t = −9.89, p < 0.001). The average reaction time (Supplementary Figure 2B) for correct trials was 1712.20 ms (SD = 475.09 ms). Pairwise comparisons showed no significant differences among any of the queried pairs (all |t|s < 3.23, all ps > 0.15) or boundary types (all |t|s < 2.50, all ps > 0.05).

### Discussion

In Experiment 2, we created a hierarchical task structure that involved lower-order (task switch) and higher-order (task and stimulus switch) event boundaries. Since only item pairs that span the higher-order boundaries involve stimuli that clearly signal distinct encoding contexts at retrieval, we predicted that order memory would be enhanced only for those pairs, but would be impaired, in line with the prior literature, for pairs spanning the lower-order boundaries. Confirming these predictions, we found that temporal order memory for pairs crossing a task switch alone was worse than for within-event stimulus pairs, but that temporal order memory for pairs crossing a task and stimulus category switch was better than for within-event stimulus pairs, which replicates Experiment 1 in demonstrating a “flip” of the classic effect.

The results for subjective temporal distance perception showed a different pattern from that for temporal order memory. Here, cross-boundary stimulus pairs were always judged to be more distant than within-event pairs, and stimulus pairs that spanned a higher-order boundary were judged as being even further apart than those that spanned a lower-order boundary.

## Experiment 3

Experiment 1 and Experiment 2 both showed that the temporal order of item pairs spanning an event boundary involving both a task and stimulus change was better remembered than that of within-event pairs. In Experiment 2, this “flip” of the classic event boundary effect on order memory was furthermore documented in the same task as the canonical effect, which was observed for boundaries that were defined by a task change only. Temporal order memory for items spanning higher-order boundaries, but not lower-order boundaries, follow the pattern of increased subjective distance ratings.

As discussed above, we hypothesized that the key factor for producing a temporal order memory advantage for cross-boundary pairs is that stimuli carry information about the encoding context at the time that order memory is being probed. In both Experiment 1 and 2, the stimuli inherently signal their context membership (e.g., the eagle in Figure 3 would inform participants that it was an object stimulus). There could be at least two mechanisms by which items evoke information about the context in which they occur. First, event features activate contexts due to pre-existing knowledge strcuctures, such as semantic memory or schemas (e.g., soapy dishes are associated with cleaning and not eating; Ghosh & Gilboa, 2014; Tulving, 1972). Second, according to the Context Maintence and Retrieval model (Polyn, 2008), event features activate their encoding context due to temporal and source associations formed during the study episode, and the context retrieved upon successful recall of an item may facilitate recollection of similarly associated items. To deconflate these two mechanisms, our experimental design in Experiment 3 orthogonalized the relationship between items of interest and their context. Finally, there could be another, non-exclusive reason, namely, that it is the magnitude or salience of the event boundary that matters, whereby a more subtle transition would disrupt temporal order memory, yet a large transition would enhance temporal order memory. Therefore, in Experiment 3, we tested whether we would continue to observe better temporal order memory for cross-boundary stimulus pairs even in a paradigm where the stimulus category remains the same and unrelated to the context throughout the experiment, as long as a context reminder is present during the memory test. To make this Experiment as comparable to the prior literature as possible, we adapted the task protocol used by Heusser and colleagues (2018) but introduced the novel manipulation of showing context cues (colored frames) at retrieval for half of the memory probes. We predicted that this context reminder would result in an order memory advantage for boundary-spanning pairs.

### Methods

#### Participants

54 participants (29 male, 25 female; ages 24~67, mean = 41.52, SD = 11.60) were recruited from MTurk for $7.00. 6 additional participants were excluded due to low accuracy (responded less than 80% of the time during the encoding phase or performed below 50% in the temporal order memory test). We doubled the target number of participants with respect to the first two experiments because in Experiment 3 we only had half the number of runs per memory probe condition (context-absent vs. context-present).

#### Stimuli

The stimulus set consisted of 468 pictures of objects from Google Image Search. The images were converted into grayscale and resized to 375 × 375 pixels on the screen. Each image was presented in conjunction with one of 10 color frames (red, orange, yellow, green, blue, purple, black, white, gray, or brown). Stimulus order and stimulus-color pairings were randomized across participants.

#### Procedure

Our experimental procedure was based on Heusser et al. (2018) and is illustrated in Figure 5. Participants performed 12 runs of the encoding and memory tasks (although due to a technical error, only the first 10 runs were recorded). During each run, participants intentionally encoded lists of 36 trial-unique grayscale objects that were presented within a colored frame. Participants were instructed to imagine the object in the color of the frame and decide if the object-color pair was aesthetically pleasing. Participants indicated their response by selecting the labels “pleasant” or “unpleasant”. The color of the frame was identical for six consecutive objects before switching to a new color for the next six objects. An event was defined as six consecutive objects with the same colored frame. There were six events per study run. On boundary trials, the frame color was updated at trial onset (i.e., concurrently with the object). Presentation timing was the same as Experiments 1 and 2.

**Figure 5.**
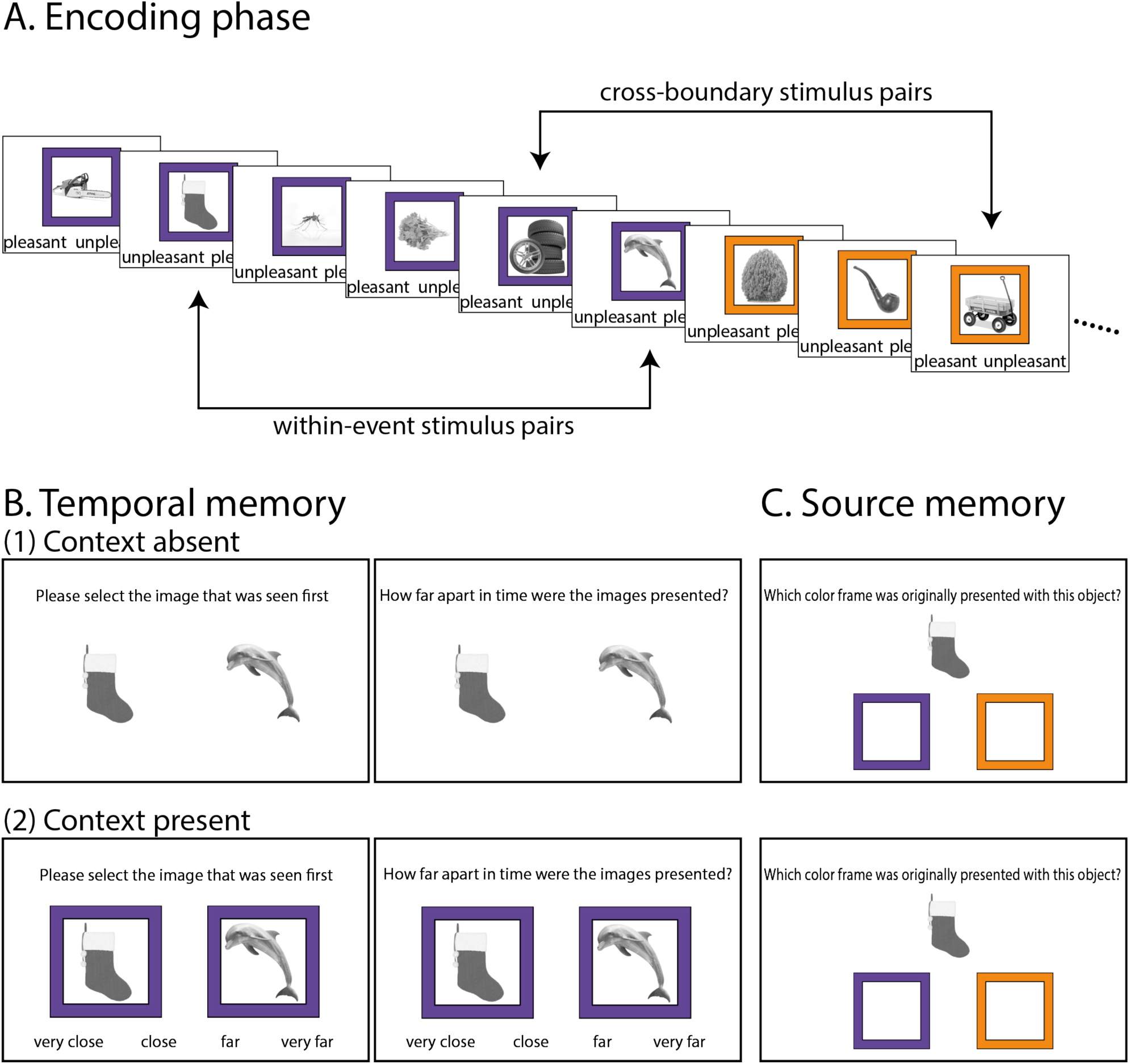
Experiment 3 & 4 design, based on Heusser et al. (2018). (A) Participants made pleasant or unpleasant judgments on object-color pairs. Critically, the color switched every six trials. (B) After encoding, participants performed a temporal memory test, consisting of a temporal order memory test (top panel) followed by a temporal distance test (bottom panel) on each trial. Half of the runs were tested with the context-absent (left panel) and half were tested with the context-present (right panel). (C) In Experiment 4 only, participants were given a source memory test in a separate block following the temporal memory test, where they were asked to indicate the colored frame originally associated with each object.

For the memory test phase, we first probed temporal order memory followed by subjective temporal distance judgment of a pair of previously studied objects. For within-event stimulus pairs we tested positions 2 and 6, and for cross-boundary stimulus pairs, we tested positions 5 and 3. There were 11 temporal memory trials in total per run – 6 within-event pairs and 5 cross-boundary pairs – presented in random order. For half of the runs, the objects were shown alone, and for the other half of the runs, the objects were shown with their colored frame. The order of context-present and context-anbsent testing runs was randomized for each participant. Crucially, participants were not told beforehand whether the colored frame would be present, and therefore could not systematically alter their encoding strategies depending on the anticipated test condition. In the practice run, half of the test trials had a colored frame, and the order was randomized across trials.

### Results

#### Encoding

Figures 6A and 6B show the encoding task mean response rates and response times for each serial position averaged across events. Response rate across all trials was high (mean = 98.16%, SD = 3.72%), with no differences across serial positions (all |t|s < 2.19, all ps > 0.27). The first trial of every new event had the longest reaction time compared to all other positions (all ts > 6.33; all ps < 0.001), reflecting a switch cost of shifting to a new frame color. Reaction times at the second trial of every event were also faster than that of the following positions (all ts > 2.98, all ps < 0.01).

**Figure 6.**
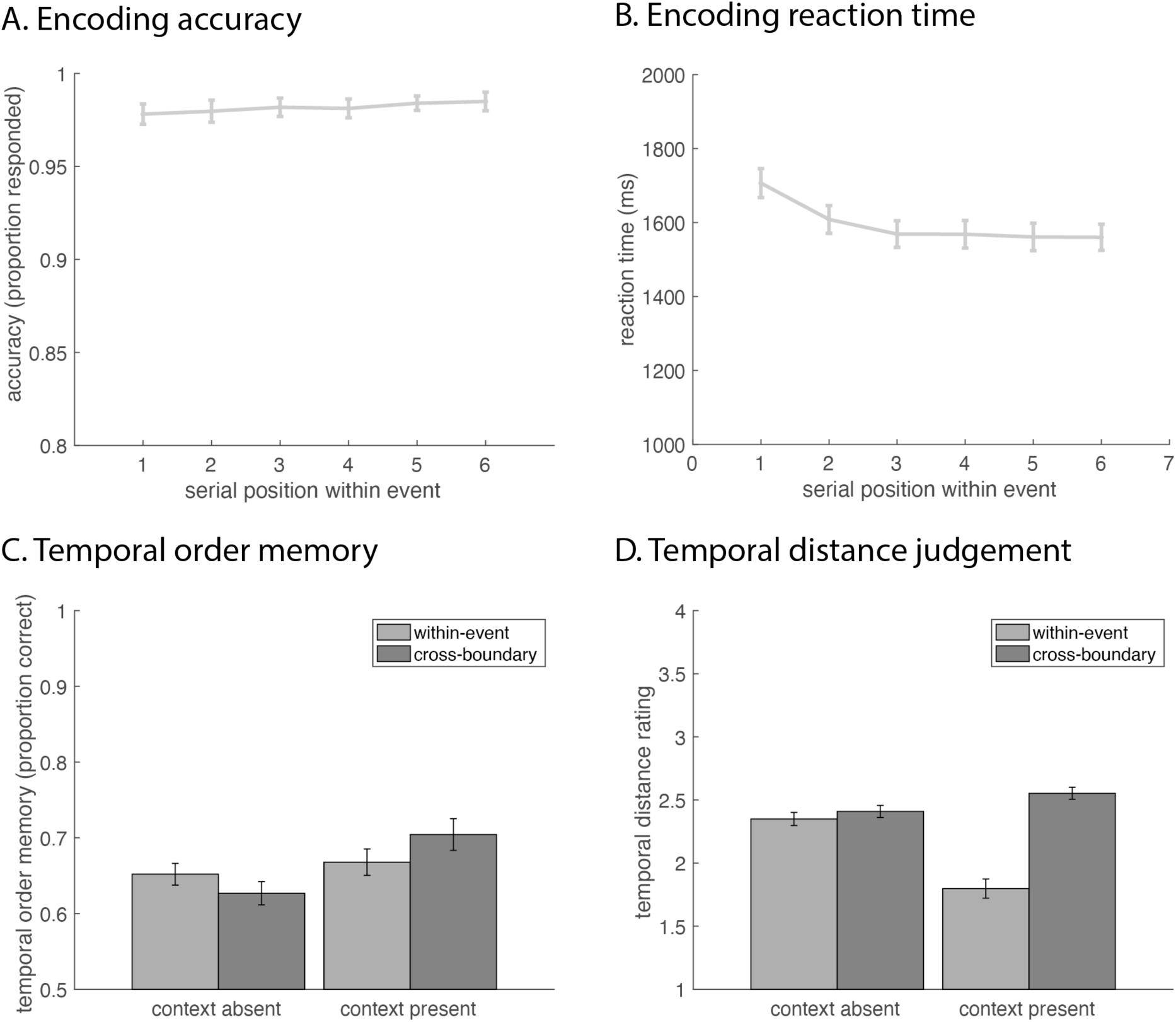
Encoding task accuracy (A) and response times (B) averaged across participants and plotted as a function of trial position within an event. Temporal order accuracy (C) and temporal distance judgment (D) for within-event and cross-boundary stimulus pairs, when the context was absent and present during test. Error bars represent standard error.

#### Temporal memory

We performed a context (absent vs. present) × boundary (within-event vs. cross-boundary) ANOVA for temporal order memory and temporal distance judgement. For temporal order memory (Figure 6C), we found a significant main effect of context (F(1,53) = 9.66, p < 0.01), with higher accuracy for runs where the context was present (mean = 68.61%, SD = 14.19) compared to when the context was absent (mean = 63.93%, SD = 10.97%) during test (t = 3.35, p = 0.001). There were no differences in order memory accuracy between within-event pairs for context-absent and context-present runs (t = −1.13, p = 0.26), however, order memory for cross-boundary pairs in the context-present runs was significantly better than in context-absent runs (t = 3.29, p < 0.01). There was no main effect of boundary (F(1,53) = 0.20, p = 0.66). Finally, there was a significant context × boundary interaction (F(1,53) = 6.34, p = 0.01). Post hoc tests showed that this interaction was driven by temporal order memory being better for cross-boundary (mean = 70.43%, SD = 15.38%) than within-event pairs (mean = 66.79%, SD = 12.76%) in the context-present condition (t = 2.02, p < 0.05), but not in the context-absent condition (t = 1.47, p = 0.15, Cohen’s d = 0.20). In the latter condition, temporal order memory was numerically better for within-events pairs (mean = 65.19%, SD = 10.57%) than cross-boundary pairs (mean = 62.67%, SD = 11.31%) for the context-absent condition, as expected. The average reaction time (Supplementary Figure 3A) for correct temporal order memory trials was 4108.42 ms (SD = 2095.91), and reaction time was not modulated by context (F(1,53) = 0.01, p = 0.91) or boundary (F(1,53) = 0.04, p = 0.84).

For temporal distance judgement (Figure 6D), we detected a significant main effect of context (F(1,53) = 33.18, p < 0.001), boundary (F(1,53) = 57.84, p < 0.001), and a context × boundary interaction (F(1,53) = 65.57, p < 0.001). Post hoc tests showed that the main effect of context was driven by temporal distance being perceived as further when the context was absent (mean = 2.38, SD = 0.37) compared to when the context was present (mean = 2.18, SD = 0.60) during retrieval (t = 6.92, p < 0.001). The main effect of boundary was driven by cross-boundary stimulus pairs (mean = 2.48, SD = 0.36) being perceived as further apart than within-event pairs (mean = 2.07, SD = 0.55; t = 6.85, p < 0.001). Finally, the interaction was driven by that cross-boundary item pairs (mean = 2.55, SD = 0.35) were perceived as further apart than within-event pairs (mean = 1.80, SD = 0.56) when the context was present (t = 8.35, p < 0.001), but not when the context was absent during retrieval (t = 1.69, p = 0.10, Cohen’s d = 0.23), although cross-boundary pairs (mean = 2.41, SD = 0.35) were numerically judged to be further apart than within-event pairs (mean = 2.35, SD = 0.38) in that condition, too. The average reaction time (Supplementary Figure 3B) for correct temporal order memory trials was 1824.82 ms (SD = 580.15), and was not modulated by context (F(1,53) = 0.07, p = 0.79) or boundary (F(1,53) = 0.15, p = 0.70), although there was a context × boundary interaction (F(1,53) = 0.07, p = 0.79) driven by faster reaction times for within-event compared to cross-boundary pairs when the context was present.

### Discussion

Experiment 3 showed that having an event context cue present at test leads to better temporal order memory for cross-boundary compared to within-event stimulus pairs, another demonstration of a “flip” of the classic finding in the event segmentation literature. While the context-absent condition matched that of previous experiments (Heusser et al., 2018), we did not find a significant effect for temporal order memory, although the pattern of results went in the same direction as in prior experiments, with within-event stimulus pairs showing numerically better temporal order memory.

With regards to temporal distance judgment, we found that when presenting the encoding context during the memory test, distance for stimulus pairs spanning an event boundary was judged to be significantly further than for pairs within the same event. There were no differences between within-event and cross-boundary pairs when the context was absent, although it showed the same trend of cross-boundary pairs being perceived as further apart.

It is important to note that there was no significant effect of context condition on reaction times in the temporal memory tests. In theory, when the context was present during the memory test, participants who remember the encoding context order could make temporal order and distance judgements based on the context alone, without having to process the individual items. Using Figure 5 as an example, if participants remembered that purple frames came before orange frames, they could correctly answer that the tires were shown before the wagon, without having to even look at the items in the context-present condition. If it were the case that participants were mainly relying on context information to make temporal judgements, we would expect reaction times to be faster in the context-present condition than the context-absent condition. However, we did not observe this effect. While a null effect should be interpreted with caution, this certainly does not speak in favor of the idea that the results of Experiment 3 were driven by context memory alone.

Taken together, the results of Experiment 3 provide additional support for the notion that event context information at retrieval enhances perceived cross-boundary item distance and leads to superior order memory for cross-boundary than within-event stimuli. Here, we deliberately preempted potential effects of schemas by using a single item category paired with unrelated contexts thoughout the experiment, therefore suggesting that schemas are not necessary for the “flip” of the classic temporal order memory effect. Additionally, since Experiment 3 only employed rather subtle task changes as event boundaries (changing color frames), this data pattern suggests that the results of Experiments 1 and 2 were not primarily driven by the magnitude or salience of event transitions, but rather by the fact that task stimuli signaled their event context membership during the memory test. The fact that in Experiment 3, participants were not told before the memory test phase whether the context would be present or not makes it unlikely that these results were driven by different encoding strategies between the context-present and context-absent runs. Since our results suggest that retrieval context affects temporal memory performance, in Experiment 4 we sought to replicate the results of Experiment 3 and to also evaluate the role of source (context) memory in potentially meditating the effects we observed.

## Experiment 4

Experiment 3 demonstrated that context information at retrieval contributes to differences in temporal order memory between within-event and cross-boundary stimulus pairs. This could be due to several reasons. For example, when the context was absent, incorrect responses could reflect source confusion, or the inability to remember the correct association of the colored frame with the object. Conversely, when the context was present at retrieval, it may give people an additional memory cue for cross-boundary pairs (since the two stimuli have different contexts), compared to the within-boundary pairs. Specifically, if one remembers the order of the contexts, then knowledge of two stimuli’s encoding source (context) at retrieval would enable one to infer their relative positions in time for cross-boundary pairs. This would suggest that superior order memory for cross-boundary item pairs should be directly related to the fidelity of people’s source memory.

In Experiment 4, we replicated the paradigm of Experiment 3 but using a between-group design. This allowed us to prevent any potential unwanted encoding strategy effects due to the mixing of context-absent and context-present retrieval conditions, as well as to obtain more data per subject for each condition. Additionally, we assessed source memory of the tested stimuli after the temporal memory phase. We hypothesized that if remembering the context associated with each stimulus were to benefit cross-boundary temporal order memory performance, this effect should be stronger for those queried stimulus pairs where both contexts were correctly remembered. Moreover, this effect should be more prominent in the context-absent condition, as here there are no external reminders of the context during retrieval, and therefore source memory should be the main determinant of whether people can use context order knowledge to inform item order judgments.

### Participants

A new set of 59 participants (27 male, 32 female; ages 23~71, mean = 41.81, SD = 12.33) was recruited from MTurk for $8.50. 6 additional participants were excluded due to low accuracy (responded less than 80% of the time during the encoding phase or performed below 50% in either the temporal order memory test or source memory test). Participants were randomly assigned to two groups, a context-absent group (N = 29) and a context-present group (N = 30), which differed only in whether the colored frame was presented during the temporal memory phase.

### Stimuli

Stimuli were the same as Experiment 3.

### Procedure

The experimental procedure was similar to Experiment 3. Participants performed 10 runs, in each of which they intentionally encoded lists of 36 trial-unique grayscale objects that were embedded in a colored frame, with the frame color changing after every six trials. After the encoding of each list, and the filler task, participants first performed 11 trials of the temporal memory task (where each trial consists of judging the temporal order of two objects, followed by rating their temporal distance). This was followed by 22 trials of the source memory task on the objects that were tested in the temporal memory task. On each test trial, participants were shown the grayscale object presented above two colored frames that were positioned on the left and right sides of the computer screen. One of these colors (target) was originally paired with the object while the other color (lure) was always one of the colored frames that had immediately preceded or followed the target color at encoding. The lure was randomized such that it was equally likely to precede or follow the target color. Participants were to indicate which colored frame originally appeared with the object.

### Results

#### Encoding

Figure 7A shows the encoding task mean response rate for each serial order position, averaged across events for the two groups. Response rate across all trials was high (mean = 97.49%, SD = 4.10%), and a context (absent vs. present) × serial position (1, 2, 3, 4, 5, and 6) ANOVA revealed no main effect of group (F(1,57) = 0.79, p = 0.38), a main effect of serial position (F(1,57) = 13.67, p < 0.001), and no group × serial position interaction (F(1,57) = 1.06, p = 0.38). We therefore combined the two groups to examine potential serial position effects. Pairwise comparisons between each serial position showed that the first trial of a new event had the lowest response rate compared to trials in all the remaining positions (all ts < −3.55, all ps < 0.01), and the second trial had a lower response rate than the fourth and fifth positions (both ts < −3.24, both ps < 0.01). Figure 7B shows the mean encoding task reaction times for each serial order position, averaged across events. A context (absent vs. present) × serial position (1, 2, 3, 4, 5, and 6) ANOVA revealed no main effect of group (F(1,57) = 0.17, p = 0.68), a main effect of serial position (F(1,57) = 57.40, p < 0.001), and no group × serial position interaction (F(1,57) = 2.01, p = 0.08). We therefore combined the two groups to examine potential serial position effects. Pairwise comparisons between each serial position showed that the first trial of a new event had the longest reaction time compared to trials in all the other positions (all ts > 8.45; all ps < 0.001). Reaction times at the second trial were also slower than the following positions (all ts > 2.60, all ps < 0.02).

**Figure 7.**
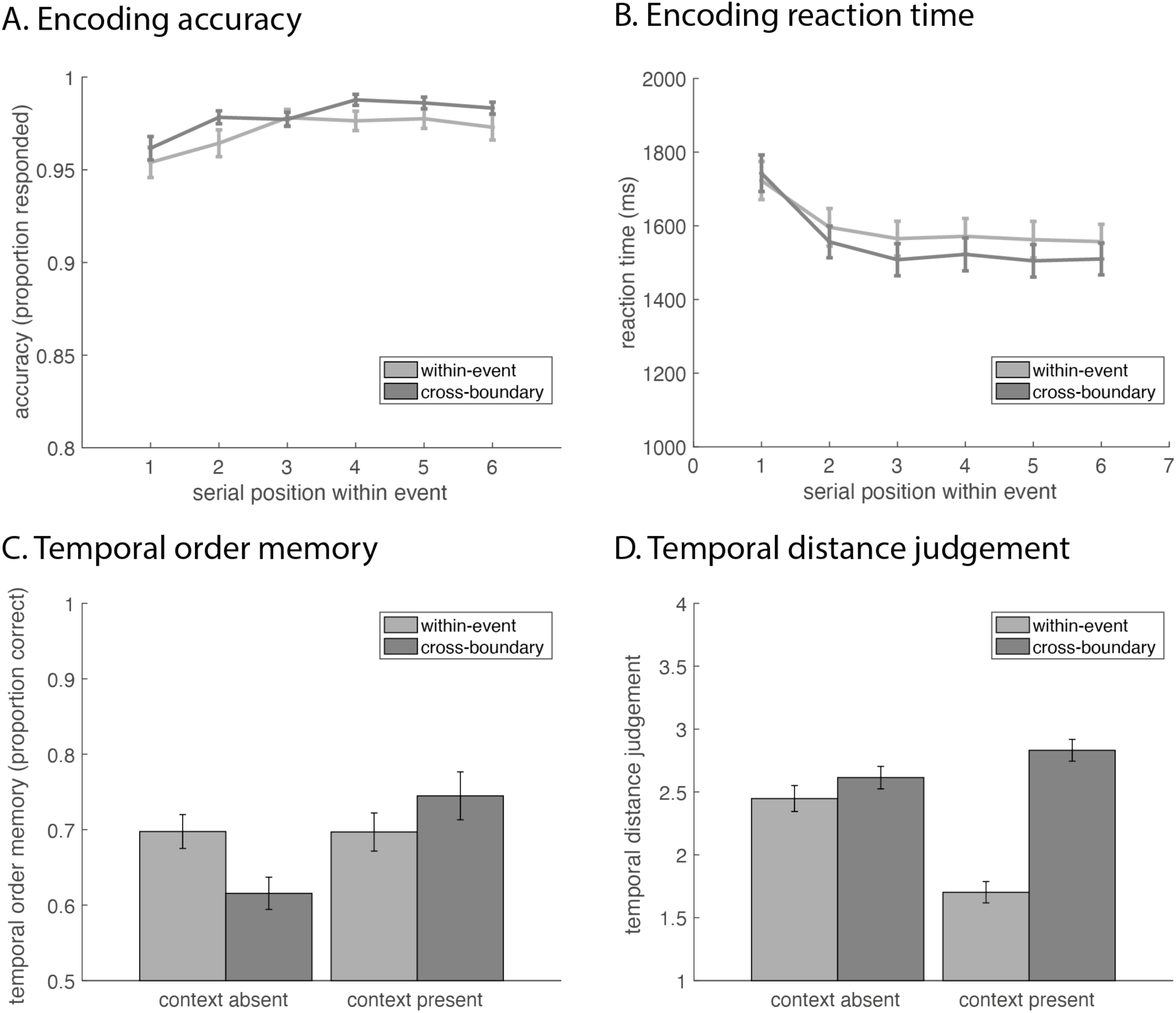
Experiment 4 encoding task accuracy (A) and response times (B) averaged across participants and plotted as a function of trial position within an event. Temporal order accuracy (C) and temporal distance judgement (D) for within-event and cross-boundary stimulus pairs, when the context was absent and present during test.

#### Temporal memory

We performed a context (absent vs. present) × boundary (within-event vs. cross-boundary) ANOVA for temporal order memory and temporal distance judgement. Figure 7C illustrates the results for temporal order memory. We found a significant main effect of context (F(1,57) = 4.15, p < 0.05). This was driven by the context-present group (mean = 72.09%, SD = 15.74%) having better temporal order memory than the context-absent group (mean = 65.66%, SD = 12.43%; t = 2.46 p = 0.02). There was no difference in order memory between within-event pairs for context-absent and context-present groups (t = 0.02, p = 0.99), however, performance for cross-boundary pairs was superior in the context-present group compared to the context-absent group (t = 3.36, p = 0.001). There was no main effect of boundary (F(1,57) = 0.91, p = 0.35). There was a significant context × boundary interaction (F(1,57) = 13.23, p < 0.001). The interaction was driven by the fact that the context-absent group showed better memory for within-event pairs (mean = 69.75%, SD = 12.12%) compared to cross-boundary pairs (mean = 61.57%, SD = 11.53%; t = 4.40, p < 0.001), whereas the context-present group did not show any significant difference in performance between within-event (mean = 69.70%, SD = 13.85%) and cross-boundary pairs (mean = 74.48%, SD = 17.33%; t = −1.59, p = 0.12, Cohen’s d = 0.29), although cross-boundary temporal order memory was numerically better than within-event order memory. With respect to reaction time, we found a context × boundary interaction for correct trials (Supplementary Figure 4A). Reaction times were slower for cross-boundary (mean = 5062.60 ms, SD = 2381.00 ms) compared to within-event (mean = 4460.11 ms, SD = 2330.32 ms) pairs in the context-present group (t = −4.22, p < 0.001), but no differences between cross-boundary (mean = 4006.25 ms, SD = 2127.00 ms) and within-event (mean = 3956.34 ms, SD = 2376.34 ms) pairs in the context-absent group (t = −0.36, p = 0.72).

Figure 7D illustrates the results for temporal distance judgement. We found a significant main effect of group (F(1,57) = 6.46, p = 0.01), with the context-absent group (mean = 2.53, SD = 0.52) rating the distances to be overall further apart than the context-present group (mean = 2.26, SD = 0.73; t = 2.24, p = 0.03). There was a significant main effect of boundary (F(1,57) = 72.23, p < 0.001), with cross-boundary pairs (mean = 2.72, SD = 0.48) being rated as further apart than within-event pairs (mean = 2.07, SD = 0.63; F(1,57) = 6.66, p < 0.001). And finally, there was a context × boundary interaction (F(1,57) = 39.84, p < 0.001). Post hoc analysis showed that the cross-boundary pairs were perceived as further apart than the within-event pairs in both the context-absent (t = 2.09, p < 0.05) and context-present (t = 8.79, p < 0.001) groups, but the effect was significantly stronger in the context-present group (t = 6.31, p < 0.001). In regards to reaction times (Supplementary Figure 4B), the context-present group (mean = 2531.76 ms, SD = 1433.30 ms) had longer reaction times compared to the context-present group (mean = 1909.67 ms, SD = 492.44 ms; t = −3.13, p < 0.01).

#### Source memory

The source memory results are shown in Figure 8. Here, we tested for a relationship between source memory and temporal order memory (Figure 8A). For each pair of objects that were tested for temporal order memory, source memory could fall into three categories, as participants may remember neither (11.12 ± 8.92 trials), one (37.07 ± 17.15 trials), or both (61.81 ± 24.67 trials) contexts correctly. We calculated the average within-event and cross-boundary temporal order accuracy for each of these three possible conditions and ran a context (absent vs. present) × source memory (remembered neither, remembered one, or remembered both) × boundary (within-event vs. cross-boundary) ANOVA. Results showed no main effect of context (F(1,42) = 1.82, p = 0.18), no main effect of source memory (F(2,84) = 2.80, p = 0.07), and no main effect of boundary (F(1,42) = 0.67, p = 0.42). We found an expected context × boundary interaction (F(1,42) = 5.58, p = 0.02). However, there was no context × source memory interaction (F(2,84) = 0.21, P = 0.81) and no source memory × boundary interaction (F(2,84) = 0.09, p = 0.91). Finally, the context × source memory × boundary interaction was also not significant (F(2,84) < 0.01, p > 0.99).

**Figure 8.**
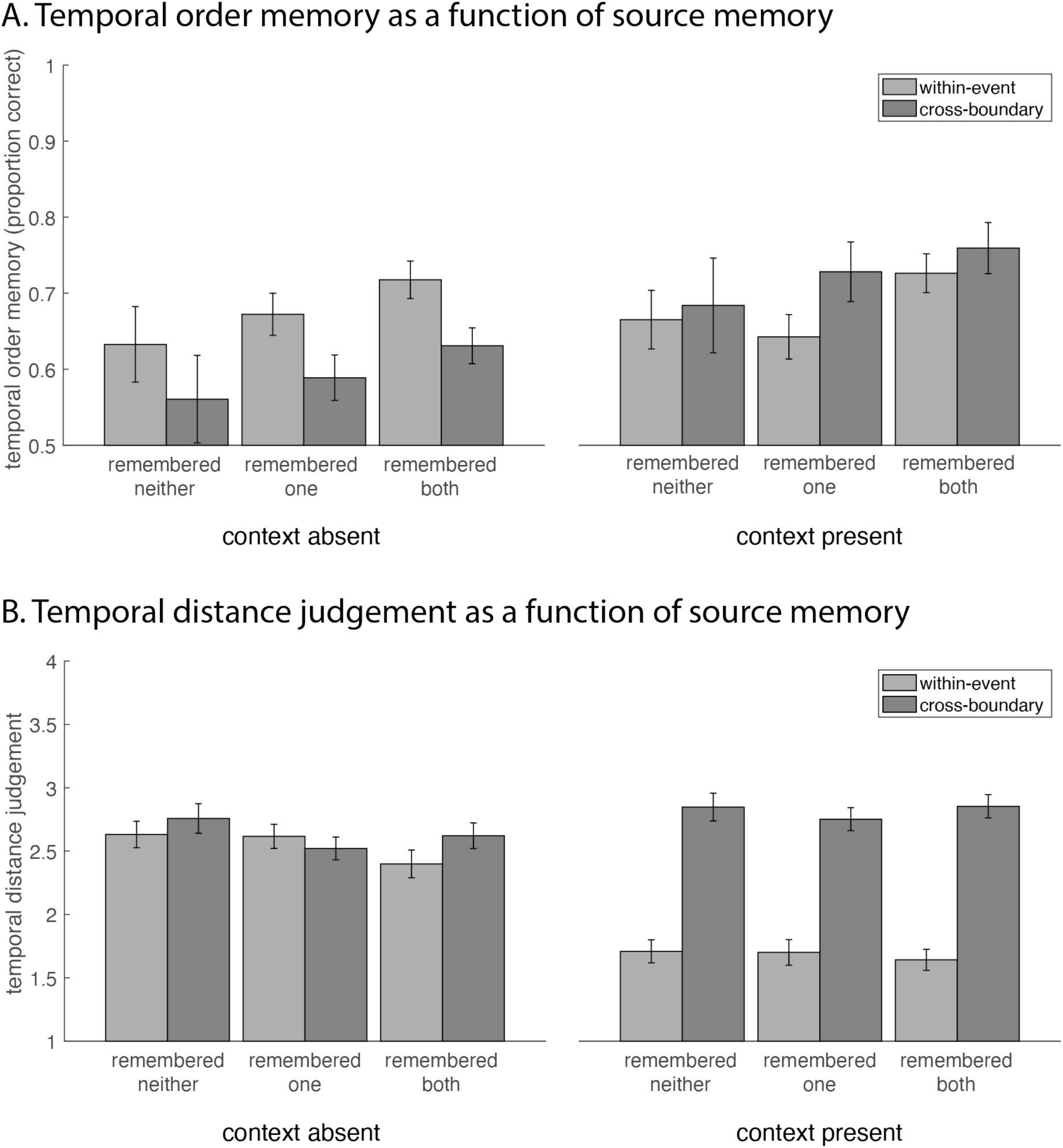
Experiment 4 temporal order memory (A) and temporal distance judgement (B) as a function of source memory for the two stimuli (remembered neither, remembered one, and remembered both). Error bars represent standard error.

We also examined the relationship between source memory and temporal distance judgement (Figure 8B), using a context (absent vs. present) × source memory (remembered neither, remembered one, or remembered both) × boundary (within-event vs. cross-boundary) ANOVA. Results showed a significant main effect of context (F(1,42) = 6.03, p = 0.02), which was driven by the context-absent group giving overall higher distance ratings (t = 4.74, p < 0.001). There was also a significant main effect of source memory (F(2,84) = 3.59, p = 0.03). Pairwise comparisons showed no difference between remembered neither (mean = 2.47, SD = 0.73) and remembered one (mean = 2.40, SD = 0.65; t = 1.83, p = 0.07), or remembered one and remembered both conditions (mean = 2.38, SD = 0.69; t = 0.91, p = 0.37). However distance ratings for when the participant remembered neither were significantly higher (rated further apart) than when they remembered both contexts (t = 2.36, p = 0.02). There was a significant main effect of boundary (F(1,42) = 42.40, p < 0.001), which was driven by cross-boundary pairs being rated as further apart than within-event pairs (t = 8.99, p < 0.001). We found no context × source memory interaction (F(2,84) = 0.92, p = 0.40), but there was a significant context × boundary interaction (F(1,42) = 41.19, p < 0.001), which was driven by the effect of cross-boundary pairs being perceived as further apart than the within-event pairs being stronger in the context-present group. There was also a source memory × boundary interaction (F(2,84) = 4.33, p = 0.02). Post hoc comparisons showed that the rating difference between cross-boundary and within-event pairs was greater when participants remembered neither or remembered both sources, compared to when participants remembered only one source (both ts > 2.07, both ps < 0.05). There was no difference between when participants remembered neither or remembered both (t = −0.34, p = 0.74). Finally, there was no context × source memory × boundary interaction (F(2,84) = 0.12, p = 0.89).

### Discussion

Experiment 4 showed that the absence of context information at test was associated with better temporal order memory for within-event compared to cross-boundary stimulus pairs, consistent with the findings in Heusser et al. (2018) and the event memory literature at large. However, when the context was present during test, we observed a (descriptive) “flip” of this effect, with the temporal order memory for cross-boundary pairs being numerically higher than for within-event pairs, which is consistent with the findings of Experiment 3.

The temporal distance judgment results also replicated the findings in Experiment 3, as temporal distance for pairs crossing an event boundary was judged to be significantly greater than for pairs within the same event, and this effect was stronger when the encoding task context was present during the memory test.

Again, we found no effect of context on reaction time in the temporal order memory test. However, we did find an effect of context on reaction time in the temporal distance judgement, in particular, participants in the context-present group had slower reaction times. If participants made judgements mainly based on context information without processing individual items, we would expect the context-present group to show a faster reaction time. Our findings show the opposite effect, suggesting that participants processed both the item and context when the latter was available.

Additionally, in this experiment, we examined whether the differences between within-event and cross-boundary pairs in temporal order memory were due to participants remembering the context associated with each stimulus. We predicted that such a source memory effect should be particularly pronounced for the context-absent condition, but we did not observe this type of interaction effect. Thus, we did not find clear evidence to support the idea that source memory determined whether temporal order memory was enhanced or impaired for cross-boundary relative to within-event stimulus pairs when the encoding context was present or absent during test. However, given the modest group sizes, lower number of trials where source memory was incorrect, and the two-alternative forced choice task design (where 50% of the responses would be correct by chance), caution should be taken when interpreting the null results of source memory and temporal order memory. We therefore followed up these results with Bayesian statistics (using JASP; JASP Team, 2022) to evaluate the strength of the evidence in favor of the null hypothesis. A Bayesian model comparison of the source memory × boundary interaction found a BF_01_ value of 12.79, indicating strong support for the null hypothesis of no interaction.

For temporal distance judgement, we found that the absence of context during retrieval led to higher temporal distance ratings. Inability to recall which event the two items belonged to also increased their perceived distance. Although, the lack of source memory could be a result of not remembering the item itself, we did not test for item recognition memory in this experiment.

## General Discussion

A substantial research literature on event memory has shown that crossing event boundaries – typically created via changes in arbitrary context cues - impairs temporal order memory for, and increases subjective temporal distance between, stimuli on each side of the boundary, compared to equally spaced stimuli within an event (e.g., DuBrow & Davachi, 2013, Heusser et al., 2018; Clewett et al., 2021). This effect seems surprising in light of intuitions about memory in everyday life, where the precise order of within-event experiences often seems more difficult to recall than the order of events per se, such that the order of stimuli stemming from different events should be more easily determined. Moreover, the fact that a greater subjective temporal distance between items was accompanied by greater confusion about their order seems paradoxical and runs counter to key models of temporal memory (Friedman, 1993, 2004; Hinrichs, 1970; Hintzman, 2002). In Experiment 1, we set out to test the possibility that temporal order for cross-boundary stimulus pairs can exceed that of within-event pairs in situations that more closely approximate the distinctive nature of events and event components we often encounter in everyday life. To this end, we created discrete contexts (including categorizing images, filling in words, math, solving puzzles, etc.), and tested temporal order memory for within- and between-event pairs of items. The results provided an initial demonstration that when items were uniquely tied to their respective encoding contexts, temporal order memory for stimuli across different contexts can be enhanced relative to those from the same event.

In Experiment 2, we created a hierarchical design where changes in task rules occurred within the same stimulus categories (e.g., performing two different object classification tasks), echoing previous experiments (e.g., Heusser et al, 2018; Clewett et al., 2021, Wang & Egner, 2022), or were accompanied by changes in task stimulus categories (e.g., moving from an object to a scene classification task), following Experiment 1. We found that temporal order memory for stimuli from adjacent events of different stimulus categories was better than that for stimuli within the same event, replicating Experiment 1. However, order memory for stimuli from the same category crossing a task switch was worse than that for within-event pairs, which is consistent with the canonical finding from prior studies. We hypothesized that the reason for these divergent results (enhancement or impairment) of cross-boundary relative to within-event temporal order memory is related to whether the stimulus signals information about the encoding event context during retrieval. The way that items can signal information about their encoding context could either be from logical inference based on semantic knowledge or schemas (Graesser & Nakamura, 1982; Tulving, 1972), or from episodic item-context bindings during encoding (Polyn, Norman, & Kahana, 2008). It is likely that both mechanism are at work in Experiment 1 and 2.

In Experiment 3, we therefore manipulated the context to be unrelated to the items during encoding, and tested whether presentation of the stimulus context at test alone was enough to benefit cross-event temporal order memory. Participants were shown grayscale images surrounded by colored frames that created event boundaries (following Heusser et al., 2018). Therefore, the stimulus category remained the same throughout the task, and event boundaries were relatively subtle. Crucially, we manipulated whether the colored frames were present or absent during the memory test. We found that temporal order memory for the within-event pairs did not differ between the context-absent and context-present conditions. However, relative to the within-event pairs, memory for cross-boundary stimulus pairs was enhanced when the context was present, and impaired when the context was absent during the temporal order memory test. In Experiment 4, we replicated this data pattern with a between-subject design and tested whether these temporal order memory effects depended on memory for the stimulus-context associations. However, our results suggested that the context-dependent temporal order memory effect was not due to an inability to retrieve the associated context. Taken together, results across the four experiments show that seemingly paradoxical predictions for cross-boundary temporal order memory based on the event memory literature versus real-life experience may be reconciled by taking into account whether encoding context information is available during retrieval. We next discuss some implications of these findings.

With respect to theories of temporal order memory, the full pattern of results in the present study cannot easily be explained by either chaining theories or distance theories. According to the chaining theory, context shifts produce a disruption to associations between stimuli, causing worse temporal order memory (DuBrow & Davachi, 2013; Gurguryan et al., 2021). By contrast, we here document several instances where event boundaries led to better rather than worse order memory (Experiments 1-4). Our results also cannot be fully explained by distance theories (Friedman, 1993, 2004; Hintzman, 2002), which posit that temporal order should be easier to discriminate the further apart in time two items are encountered or perceived (Rouhani et al., 2020). However, despite the fact that participants rated temporal distance to increase between stimuli crossing an event boundary (Ezzyat & Davachi, 2014; Lositsky et al., 2016; Sahakyan & Smith, 2014), we documented impaired temporal order memory for cross-boundary compared to within-event stimulus pairs when no context information was available during retrieval (Experiments 2-4). We additionally show in Experiment 4, that the direction of enhanced or impaired temporal order memory seems to be independent of source memory. This suggests that temporal order judgment of cross-boundary stimuli is not dependent on whether one can remember the associated encoding context, but if that context is externally provided during retrieval, it clearly enhances cross-boundary order memory. One way to reconcile these two observations could be that people may not typically attempt to actively retrieve the encoding context unless prompted (in the source memory test). Another possibility is that when the encoding context is absent, retrieving the source information may lead to interference or confusion of the order of cross-boundary items during test by bringing temporally segregated events to mind simultaneously.

In particular, similar to our pattern of results Cox et al. (2021) found that whether context was reinstated at retrieval could impair or enhance associative memory. In their study, participants learned to associate word pairs (AB) on unique context background images on Day 1. On Day 2, participants learned new word pairs (AC) depicted either against the same background contexts as Day 1 or against new contexts. Finally, on Day 3, participants performed cued recall tests. During test, if the original context was absent, AC learning in the same context markedly enhanced the original (AB), new (AC), and inferential (BC) memories. However, this difference disappeared when the context was present during test. The authors proposed that items studied in the same context elicit integration of memories during learning (Schlichting & Preston, 2014; Wahlheim et al., 2015), but that contextual reinstatement during recall leads to interference. Conversely, context changes across episodes induce competition between memories, thus promoting new learning at the cost of the original memories (Radvansky & Copeland, 2006), but these effects can be overturned by contextual reinstatement during retrieval. This conjecture suggests that memory disruption caused by event boundaries is not permanent, resembling the effects observed in context-dependent memory (Godden & Baddeley, 1975; Smith, 1979; Smith & Vela, 2001). For example, one can imagine that in a scenario of moving from the UK to the US (context change), it would be functional to update memory for new stimuli at the expense of old memories, e.g., remember that “chips” are called “fries”. However, such retroactive forgetting should not be permanent, and the correct term should be the one most easily retrieved in the country it is associated with. Similarly, memories for these stimuli when the original context is absent, for example, if you were teaching English in a third, non-English speaking country, may compete with each other, such that “chips” and “fries” would produce retrieval interference (Anderson, 2003; Ritvo, Turk-Browne, & Norman, 2019; Wimber, Alink, Charest, Kriegeskorte, & Anderson, 2015). It is possible that similar context-dependent mechanisms may be involved during the retrieval of temporal order memory.

It has previously been shown that schematic event structure created by naturalistic event boundaries can be exploited for remembering event components (Boltz, 1992; Brewer & Dupree, 1983; Lichtenstein & Brewer, 1980). In real life, events themselves often follow a certain chronoligical order (Schank & Abelson, 1977). For example, dishes would only require cleaning after one has eaten. Thus, if queried about the temporal order of these two events, one could rely on merely reasoning that the clean-up should happen after eating, without having to engage in episodic memory recall. In our Experiments 1-4, contexts were arbitrarily generated and chronologically unrelated, such that participants would need to engage in memory recall. While it is theoretically possible that in the context-present conditions, our participants could rely only on order memory for the contexts during the memory tests without processing the individual items, the reaction time analysis in Experiments 3 and 4 suggested that participants processed both the item and context in our experiments. For future studies, it may be interesting to examine the effect of event boundaries using events with an inherent chronological order. This could be conducted using more complex, everyday events as stimulus material, such as movies or narratives. Alternatively, one could examine temporal order memory as a function of learning using a fixed order of events across runs.

Furthermore, unlike in most event boundary studies, real-life event perception and task goals are typically hierarchical, with smaller events or steps nested within larger overarching ones (Badre & Nee, 2018; Baldassano et al., 2017; Hasson, Yang, Vallines, Heeger, & Rubin, 2008; Hasson, Chen, & Honey, 2015; Speer, Zacks, & Reynolds, 2007). For example, one may experience two task episodes, “eat breakfast” and “clean dishes”, in one morning. Each episode may be further broken down into smaller events, such as “open fridge… slice bagel… spread cream cheese… put on plate” and “turn on tap… squeeze soap… scrub plate… dry with towel” (Cooper & Shallice, 2000; Schneider & Logan, 2006; Wen, Duncan, & Mitchell, 2020). At the topmost level of the hierarchy, representations are at a temporally and conceptually broader scale and have been discussed in terms of schemas (Robin & Moscovitch, 2017), event models (Stawarczyk, Bezdek, & Zacks, 2021), and situation models (Reagh & Ranganath, 2018). In real life, context therefore typically reflects slowly drifting information and associated schemas can be used to organize our memories of more transient information (Manning et al., 2013). Neural representation of event boundaries at the higher, schematic level have been found to involve different brain regions than lower-order, and more transient task transitions, with the former involving the default mode network, and the latter being coded in the multiple demand network (Badre & Nee, 2018; Crittenden et al., 2015; Wen et al., 2020). These previous studies suggest that event segmentation occurs at multiple time scales that directly correspond to the magnitude and meaningfulness of event transitions (Baldassano et al., 2017). While in the current study, the temporal memory effects observed in Experiments 1 and 2 seem stronger than in Experiments 3 and 4, which would tentatively suggest that context distinctiveness may play a role in event boundaries (Gurguryan et al., 2021), we cannot directly compare the Experiments due to the different number of events and trials within each experiment. However, examining the hierarical organization of schemas and its effect on temporal order memory would be an interesting avenue for future studies.

Finally, in Experiments 1 and 2, we pursued a novel testing strategy in this literature by probing memory for stimulus pairs that cover the entire range of each context rather than focusing only on a single pre-selected within-event and cross-boundary pair. This allowed us to establish whether there were any serial position effects (Henson, 1998; Hintzman, Block, & Summers, 1973) on temporal memory. We found enhancement of temporal order memory for stimuli at the beginning of a new context, and that accuracy decreased with serial position within an event. Participants also rated the stimulus pairs containing the first stimuli of an event to be more distant than other within-event pairs. Prior studies have reported better overall memory for information studied at event boundaries than information at non-boundary positions (Heusser et al., 2018; Pettijohn, Thompson, Tamplin, Krawietz, & Radvansky, 2016; Polyn, Norman, & Kahana, 2009; Swallow, Zacks, & Abrams, 2009), including better item memory and item-context associations. This is possibly due to segregation of information into separate event models decreasing retroactive interference (Bower, Clark, Lesgold, & Winzenz, 1969; Smith, Glenberg, & Bjork, 1978; Strand, 1970). Other studies on contextual novelty have found that a context shift during encoding creates a mismatch between the new task representation and the previous task encoding representation, triggering more focused attention on the first post-shift item, which gives it a boost in memory (Fabiani & Donchin, 1995; Köhler & von Restorff, 1933; Lin, Pype, Murray, & Boynton, 2010; Ranganath & Rainer, 2003). In a related study, Polyn et al. (2009) showed that switching encoding tasks midway while encoding a short list of words resulted in a significant increase in the free recall of words that followed the task switch. In our experiments, the beginning of each new task context would introduce such novelty. Consequently, we showed that the temporal order of both early within-event pairs and cross-boundary pairs was better remembered. In a recent study, Pu et al. (2022) also found a local primacy effect using varying event lengths. The authors proposed a computational model where event boundaries cause a systematic change in temporal context by reinstating a certain proportion of the very first contextual representation, which accounts for both the local primacy effect as well as the classic event boundary effect. Finally, it is also possible that the salient event boundaries created in experiments 1 and 2 could have served as a “tag” or “landmark” signaling the start or end of an event (Henson, 1998). Previous studies have found that landmark events, such as the eruption of Mt. St. Helens, an earthquake, New Year’s Day, and even one’s own personal landmarks improved the accuracy of recalling when a memory occurred (Friedman, 2004; Loftus & Marburger, 1983). Landmarks are useful, as one can refer back to a salient stimulus or transition to infer the timing of surrounding events. In line with the landmark hypothesis, Michelmann and colleagues (2021; 2019) proposed that event boundaries serve as access points during continuous memory retrieval, such that when scanning a memory for a target, people can skip ahread to the beginning of new events, speeding up memory scanning time. Taken together, our results suggest that event perception and memory are dependent on the stimuli’s positions within events and can involve multiple cognitive processes.

In conclusion, the present study demonstrates a retrieval context-dependence of event boundary effects on memory, whereby the presence of encoding context information at retrieval enhances temporal order memory for stimuli spanning an event boundary relative to order memory for within-event stimuli. The presence or absence of schema or context information at retrieval may cue people to focus either on the order of broader episodes or mentally time-travel within an event, respectively, and thereby support the flexible nature of episodic memory.

