## Supplementary for "Retrieval context determines whether event boundaries impair or enhance temporal order memory"

**Supplementary Material**

**Experiment 1 Memory Test Reaction Times**

Supplementary Figure 1A illustrates reaction times during the temporal order memory test as a function of serial order during encoding. Pairwise comparisons showed that among within-events pairs, the first pair, (1 vs. 4), had significantly faster reaction times than pairs (3 vs. 6), (5 vs. 8), (6 vs. 9), and (7 vs. 10) (all ts < -3.14, all ps < 0.04). (2 vs. 5) also had faster reaction times than (5 vs. 8) (t = -3.86, p = 0.03). Reaction time differences between all other pairs were insignificant (all ts < 2.92, all ps > 0.05). The mean reaction time for correct trials was 2570.17 ms (SD = 796.22 ms), and we found no significant differences comparing the average reaction times for within-event and cross-boundary pairs (t = 1.67, p = 0.11).

Supplementary Figure 1B illustrates reaction times of the temporal distance judgement task as a function of serial order during encoding. Pairwise comparisons showed significant differences between numerous queried pairs. Most prominently, cross-boundary pair (8 vs. 1) had slower reaction times than within-event pairs (3 vs. 6), (4 vs. 7), (5 vs. 8), (6 vs. 9), and (7 vs. 10) (all ts < -2.85, all ps < 0.04). Cross-boundary pair (9 vs. 2) also had slower reaction times than within-event pairs (3 vs. 6), (6 vs. 9), and (7 vs. 10) (all ts < -2.74, all ps < 0.05). Combing all within-event (mean = 1617.26 ms, SD = 484.37 ms) and cross-boundary pairs (mean = 1888.89 ms, SD = 657.22 ms), we found that cross-boundary pairs overall slower reaction times (t = -2.85, p < 0.01).

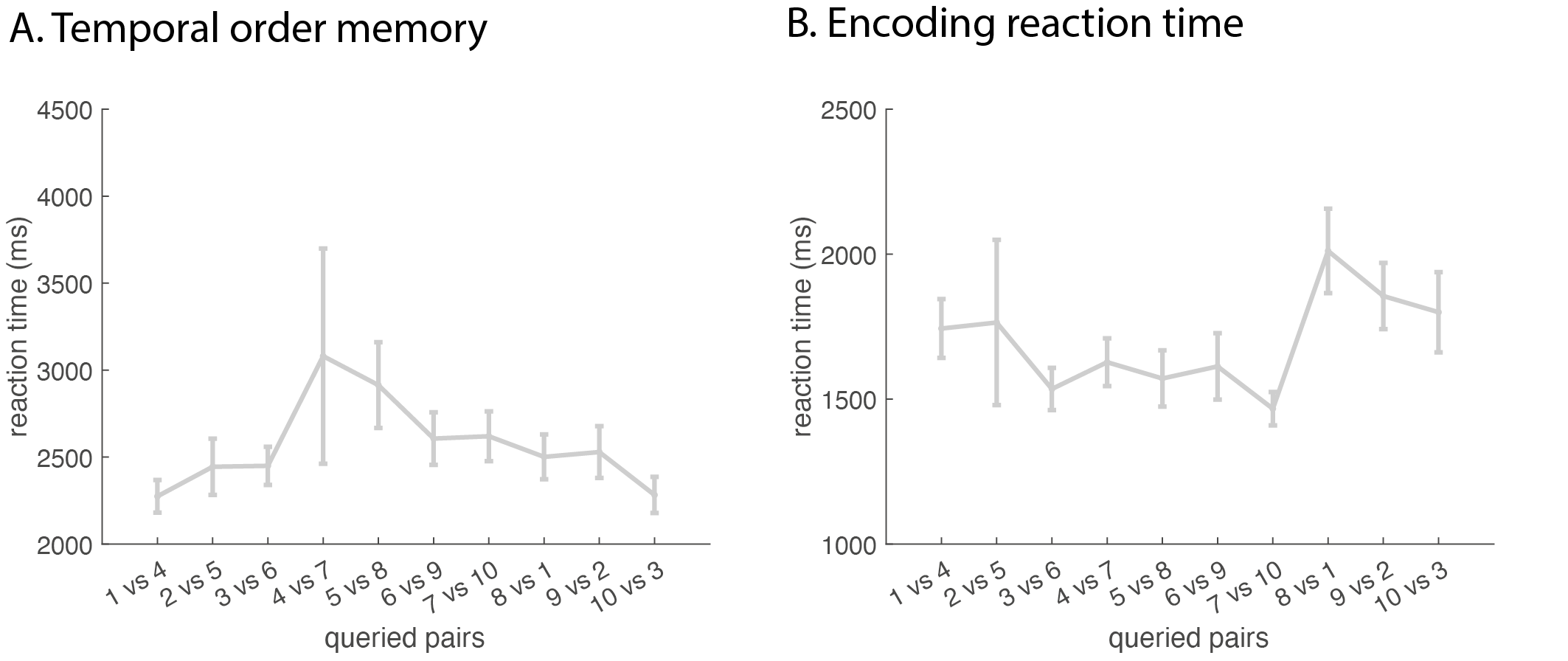

*Supplementary Figure 1.* *Experiment 1 reaction times of the temporal order memory test (A) and temporal distance judgement (B) averaged across participants and plotted as a function of trial position pairs. Error bars represent standard error.*

**Experiment 2 Memory Test Reaction Times**

Supplementary Figure 2A illustrates reaction times of the temporal order memory test as a function of serial position and boundary type. The only significant differences that emerged from pairwise comparisons were slower reaction times for (5 vs. 8) compared to (8 vs. 1) (t = -4.31, p < 0.01). Comparisons among item pairs grouped according to boundary type showed that participants were faster at recalling the temporal order of stimuli spanning a higher-order boundary (mean = 2388.80 ms, SD = 1043.45 ms) compared to stimuli within the same event (mean = 2688.85 ms, SD = 1218.05 ms; t = 3.42, p < 0.01) as well as those spanning a lower-order boundary (mean = 2792.67 ms, SD = 1109.84 ms; t = 3.00, p < 0.01). There were no differences in reaction time for stimuli within the same event and those spanning a lower-order boundary (t = -0.93, p = 0.36).

Supplementary Figure 2B illustrates reaction times of the temporal distance judgement task as a function of serial position and boundary type. The average reaction time for correct trials was 1712.20 ms (SD = 475.09 ms). Pairwise comparisons showed no significant differences among any of the queried pairs (all |t|s < 3.23, all ps > 0.15). Furthermore, there were no significant differences among boundary types after combining trials of the same type (all |t|s < 2.50, all ps > 0.05).

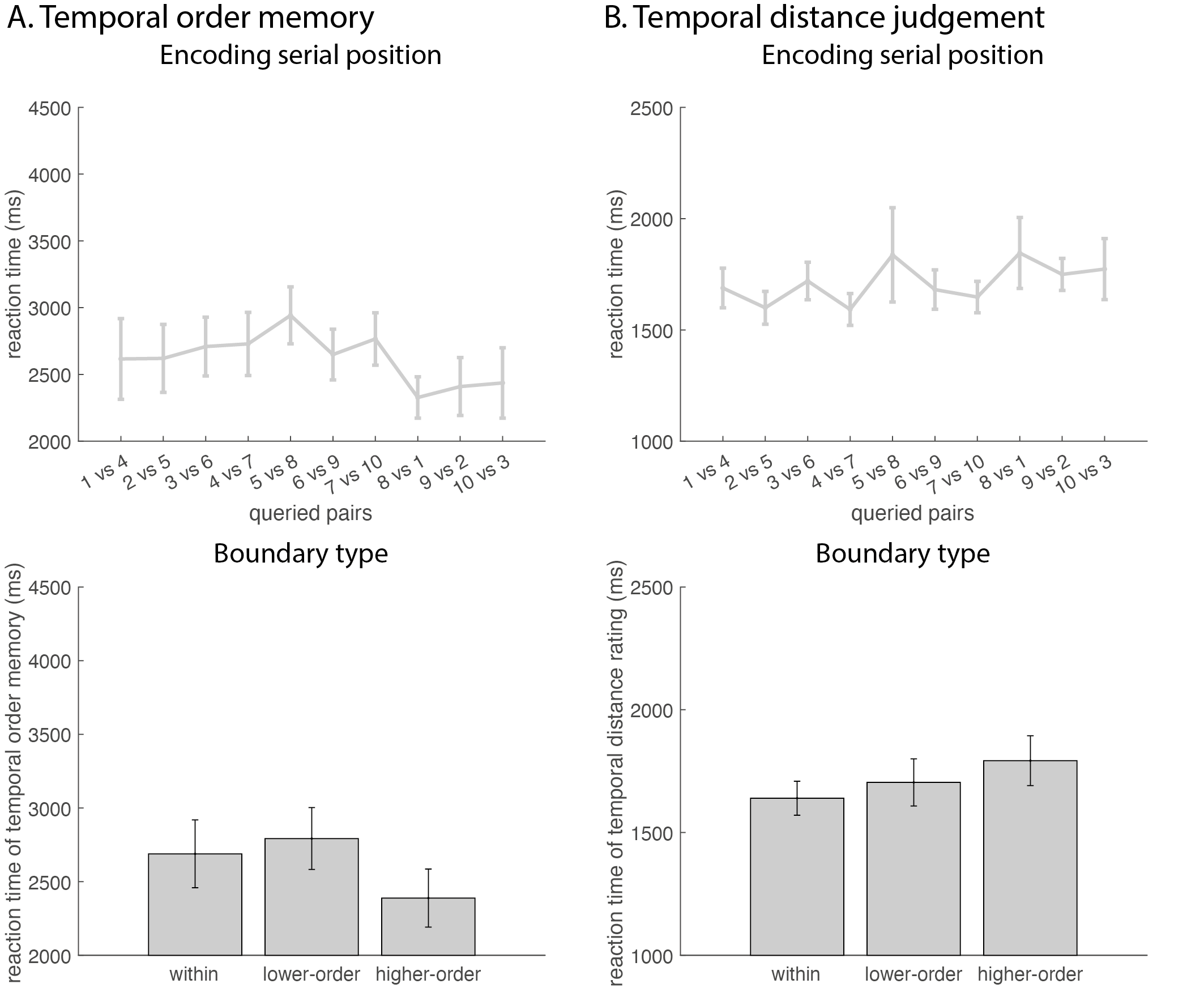

*Supplementary Figure 2.* *Experiment 2 reaction times of the temporal order memory test (A) and temporal distance judgement (B) averaged across participants and plotted as a function of trial position pairs (top) and boundary type (bottom; within-event, task switch, and task and context switch). Error bars represent standard error.*

**Experiment 3 Memory Test Reaction Times**

Supplementary Figure 3A illustrates reaction times during the temporal order memory test (mean = 4108.42 ms, SD = 2095.91). A context (absent vs. present) × boundary (within-event vs. cross-boundary) ANOVA showed no main effect of context (F(1,53) = 0.01, p = 0.91), no main effect of boundary (F(1,53) = 0.04, p = 0.84), and no context × boundary interaction (F(1,53) = 1.95, p = 0.17).

Supplementary Figure 3B illustrates reaction times during the temporal distance judgement (mean = 1824.82 ms, SD = 580.15). A context (absent vs. present) × boundary (within-event vs. cross-boundary) ANOVA showed no main effect of context (F(1,53) = 0.07, p = 0.79) and no main effect of boundary (F(1,53) = 0.15, p = 0.70). There was a significant context × boundary interaction (F(1,53) = 7.76, p < 0.01). Post hoc analysis showed that this interaction was driven by no reaction time differences between the within-event and cross-boundary pairs when the context was absent (t = 1.95, p = 0.06), but faster reaction times for within-event pairs when the context was present (t = -2.22, p = 0.03).

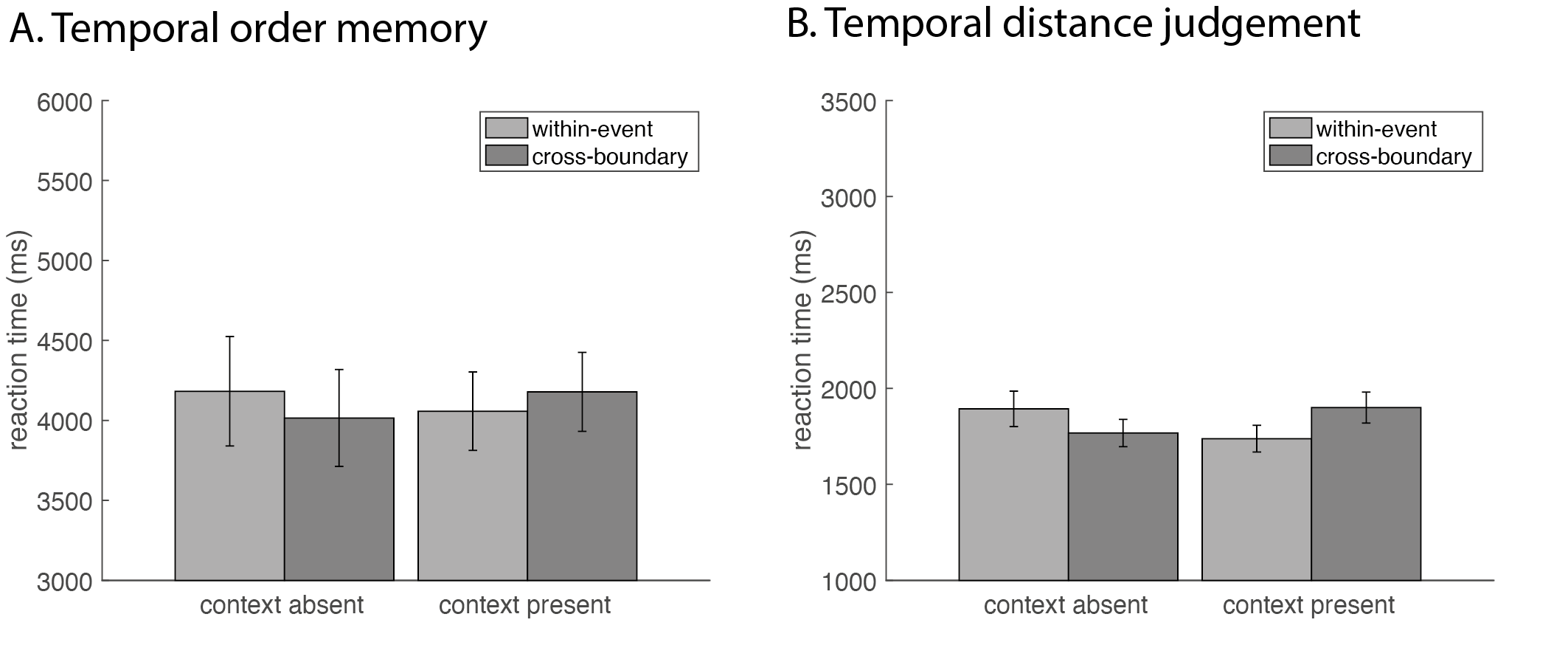

*Supplementary Figure 3.* *Experiment 3 reaction times during the temporal order memory test (A) and temporal distance judgement (B) for within-event and cross-boundary stimulus pairs, when the context was absent and present during test. Error bars represent standard error.*

**Experiment 4 Memory Test Reaction Times**

Supplementary Figure 4A illustrates reaction times during the temporal order memory test. A context (absent vs. present) × boundary (within-event vs. cross-boundary) ANOVA showed no main effect of context (F(1,57) = 1.73, p = 0.19). There was a significant main effect of boundary (F(1,57) = 10.66, p < 0.01) and a context × boundary interaction effect (F(1,57) = 7.65, p < 0.01). Post hoc analysis showed that the effects were driven by a slower reaction time for cross-boundary (mean = 5062.60 ms, SD = 2381.00 ms) compared to within-event (mean = 4460.11 ms, SD = 2330.32 ms) pairs in the context-present group (t = -4.22, p < 0.001), but no differences between cross-boundary (mean = 4006.25 ms, SD = 2127.00 ms) and within-event (mean = 3956.34 ms, SD = 2376.34 ms) pairs in the context-absent group (t = -0.36, p = 0.72).

Supplementary Figure 4B illustrates reaction times for the temporal distance judgement. A context (absent vs. present) × boundary (within-event vs. cross-boundary) ANOVA showed a main effect of context (F(1,57) = 5.86, p = 0.02). This was driven by the context-present group (mean = 2531.76 ms, SD = 1433.30 ms) having overall slower reaction times compared to the context-absent group (mean = 1909.67 ms, SD = 492.44 ms; t = -3.13, p < 0.01). There was no main effect of boundary (F(1,57) = 2.01, p = 0.16) and no context × boundary interaction (F(1,57) = 2.06, p = 0.16).

Supplementary Figure 4C plots the reaction times during the temporal order memory test as a function of source memory for the two stimuli (remembered neither, remembered one, and remembered both). We calculated the average reaction times for within-event and cross-boundary temporal order memory for each of the three source memory conditions and ran a context (absent vs. present) × source memory (remembered neither, remembered one, or remembered both) × boundary (within-event vs. cross-boundary) ANOVA. Results showed no main effect of context (F(1,42) = 1.13, p = 0.29), no main effect of source memory (F(2,84) = 1.82, p = 0.17), and no main effect of boundary (F(1,42) = 1.21, p = 0.28). We found a significant context × boundary interaction (F(1,42) = 6.52, p = 0.01). This interaction was driven by the fact that while there were no reaction time differences between within-event versus cross-boundary item pairs in the context-absent group (t = 0.43, p = 0.67), in the context-present group, cross-boundary pairs had a slower reaction time than within-event pairs (t = -4.32, p < 0.001). There was no context × source memory interaction (F(2,84) = 1.27, P = 0.29) and no source memory × boundary interaction (F(2,84) = 0.18, p = 0.83). The context × source memory × boundary interaction was also not significant (F(2,84) = 1.97, p = 0.15).

Supplementary Figure 4D plots the reaction times during the temporal distance judgement as a function of source memory for the two stimuli. We calculated the average reaction times for within-event and cross-boundary temporal order memory for each of the three source memory conditions and ran a context (absent vs. present) × source memory (remembered neither, remembered one, or remembered both) × boundary (within-event vs. cross-boundary) ANOVA. Results showed a main effect of context (F(1,42) = 6.02, p = 0.02), which was driven by longer reaction times in the context-present group (t = 4.09, p < 0.001). There was no main effect of source memory (F(2,84) = 1.15, p = 0.32) or boundary (F(1,42) = 0.78, p = 0.38). We found a source memory × boundary interaction (F(2,84) = 3.31, p = 0.04). Pairwise comparisons showed that the difference in reaction time between judging cross-boundary and within-event pairs was smaller when participants remembered neither context compared to when the participant remembered one context (t = -2.03, p < 0.05). However, the boundary difference was not significant between remembering neither and remembering both contexts or remembering one and remembering both contexts (both ts > -1.97, both ts > 0.05). We found no context × source memory interaction (F(2,84) = 0.27, p = 0.76), no context × boundary interaction (F(1,42) = 0.24, p = 0.63), and no context × source memory × boundary interaction (F(2,84) = 1.19, p = 0.31).

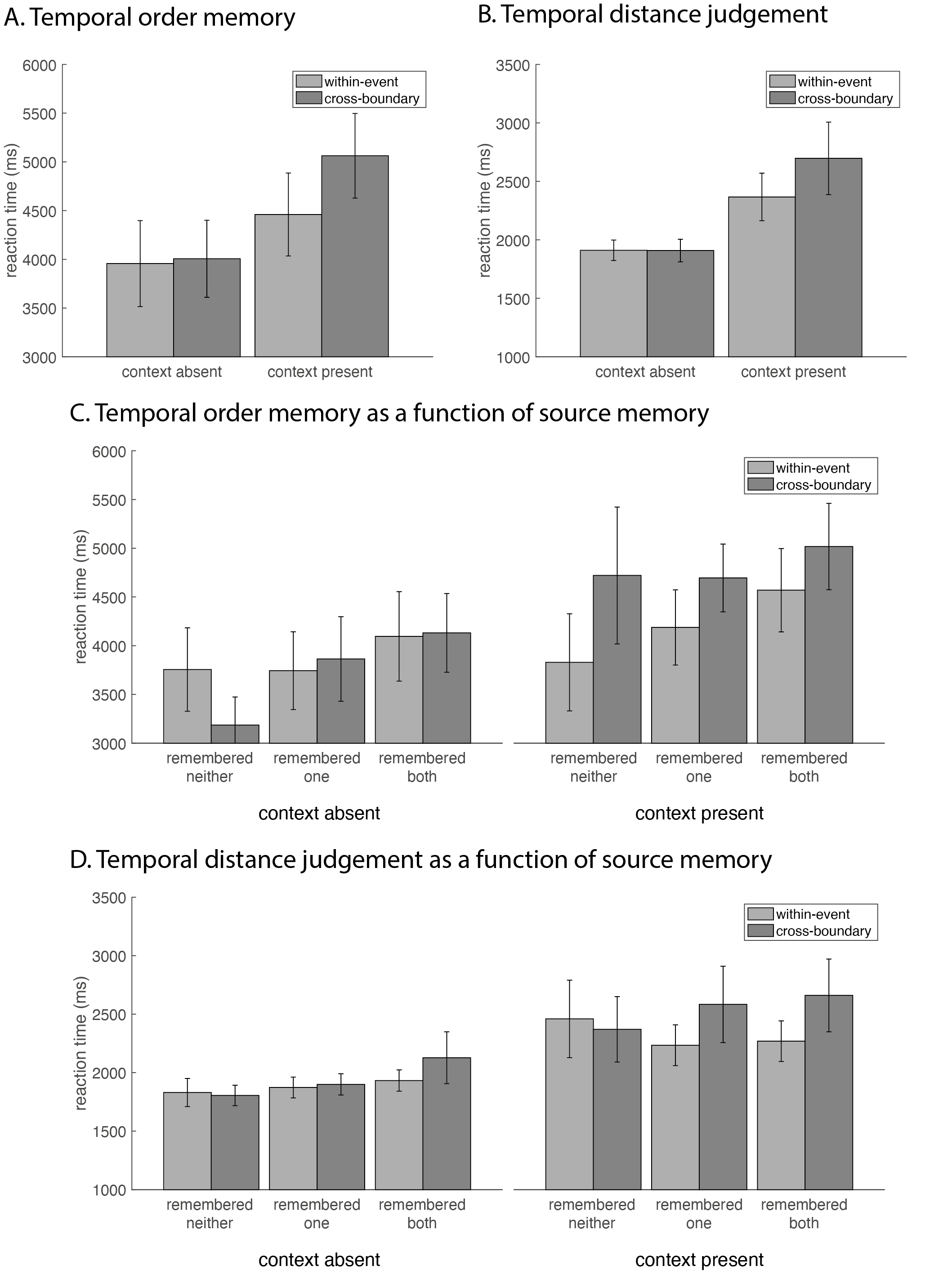

*Supplementary Figure 4.* *Experiment 4 reaction times during the temporal order memory test (A) and temporal distance judgement (B) for within-event and cross-boundary stimulus pairs, when the context was absent and present during test. (C). Reaction times during the temporal order memory test as a function of source memory for the two stimuli (remembered neither, remembered one, and remembered both). (D) (C). Reaction times during the temporal distance judgement as a function of source memory for the two stimuli. Error bars represent standard error.*

| Position | 1 vs 4 | 2 vs 5 | 3 vs 6 | 4 vs 7 | 5 vs 8 | 6 vs 9 | 7 vs 10 | 8 vs 1 | 9 vs 2 | 10 vs 3 |
| --- | --- | --- | --- | --- | --- | --- | --- | --- | --- | --- |
| 1 vs 4 |  | t = 1.93  p < 0.001 | t = 11.63  p < 0.001 | t = 9.24  p < 0.001 | t = 8.68  p < 0.001 | t = 13.19  p < 0.001 | t = 14.08  p < 0.001 | t = -0.75  p = 0.52 | t = -0.70  p = 0.54 | t = -1.56  p = 0.15 |
| 2 vs 5 |  |  | t = 9.74  p < 0.001 | t = 7.31  p < 0.001 | t = 7.36  p < 0.001 | t = 10.49  p < 0.001 | t = 11.12  p < 0.001 | t = -2.20  p < 0.05 | t = -2.05  p = 0.06 | t = -2.32  p = 0.04 |
| 3 vs 6 |  |  |  | t = 0.59  p = 0.60 | t = 2.44  p = 0.03 | t = 4.71  p < 0.001 | t = 6.34  p < 0.001 | t = -12.44  p < 0.001 | t = -8.48  p < 0.001 | t = -10.09  p < 0.001 |
| 4 vs 7 |  |  |  |  | t = 1.73  p = 0.11 | t = 3.28  p < 0.01 | t = 4.40  p < 0.001 | t = -8.64  p < 0.001 | t = -7.52  p < 0.001 | t = -9.29  p < 0.001 |
| 5 vs 8 |  |  |  |  |  | t = 1.36  p = 0.21 | t = 2.58  p = 0.02 | t = -9.63  p < 0.001 | t = -7.59  p < 0.001 | t = -7.88  p < 0.001 |
| 6 vs 9 |  |  |  |  |  |  | t = 1.86  p = 0.09 | t = -13.40  p < 0.001 | t = -10.76  p < 0.001 | t = -11.35  p < 0.001 |
| 7 vs 10 |  |  |  |  |  |  |  | t = -15.22  p < 0.001 | t = -11.76  p < 0.001 | t = -13.27  p < 0.001 |
| 8 vs 1 |  |  |  |  |  |  |  |  | t = -0.12  p = 0.90 | t = -0.52  p = 0.64 |
| 9 vs 2 |  |  |  |  |  |  |  |  |  | t = -0.36  p = 0.74 |
| 10 vs 3 |  |  |  |  |  |  |  |  |  |  |

Table S1. Comparisons between queried temporal order memory pairs in Experiment 1

Table S2. Comparisons between queried temporal distance memory pairs in Experiment 1

| Position | 1 vs 4 | 2 vs 5 | 3 vs 6 | 4 vs 7 | 5 vs 8 | 6 vs 9 | 7 vs 10 | 8 vs 1 | 9 vs 2 | 10 vs 3 |
| --- | --- | --- | --- | --- | --- | --- | --- | --- | --- | --- |
| 1 vs 4 |  | t = 11.83  p < 0.001 | t = 10.60  p < 0.001 | t = 11.71  p < 0.001 | t = 10.67  p < 0.001 | t = 10.63  p < 0.001 | t = 10.13  p < 0.001 | t = -12.47  p < 0.001 | t = -13.58  p < 0.001 | t = -15.56  p < 0.001 |
| 2 vs 5 |  |  | t = -1.22  p = 0.33 | t = -0.34  p = 0.83 | t = -0.06  p = 0.96 | t = -0.59  p = 0.72 | t = -0.52  p = 0.74 | t = -13.13  p < 0.001 | t = -13.91  p < 0.001 | t = -14.30  p < 0.001 |
| 3 vs 6 |  |  |  | t = 0.90  p = 0.53 | t = 1.48  p = 0.22 | t = 0.39  p = 0.83 | t = 0.70  p = 0.66 | t = -12.77  p < 0.001 | t = -13.36  p < 0.001 | t = -12.91  p < 0.001 |
| 4 vs 7 |  |  |  |  | t = 0.27  p = 0.87 | t = -0.34  p = 0.83 | t = -0.18  p = 0.92 | t = -13.55  p < 0.001 | t = -14.25  p < 0.001 | t = -14.67  p < 0.001 |
| 5 vs 8 |  |  |  |  |  | t = -0.59  p = 0.72 | t = -0.52  p = 0.74 | t = -12.98  p < 0.001 | t = -13.67  p < 0.001 | t = -14.01  p < 0.001 |
| 6 vs 9 |  |  |  |  |  |  | t = 0.16  p = 0.92 | t = -13.13  p < 0.001 | t = -13.77  p < 0.001 | t = -13.91  p < 0.001 |
| 7 vs 10 |  |  |  |  |  |  |  | t = -12.71  p < 0.001 | t = -13.28  p < 0.001 | t = -13.55  p < 0.001 |
| 8 vs 1 |  |  |  |  |  |  |  |  | t = -3.61  p < 0.01 | t = -2.69  p = 0.02 |
| 9 vs 2 |  |  |  |  |  |  |  |  |  | t = 0.06  p = 0.96 |
| 10 vs 3 |  |  |  |  |  |  |  |  |  |  |

Table S3. Comparisons between queried temporal order memory pairs in Experiment 2

| Position | 1 vs 4 | 2 vs 5 | 3 vs 6 | 4 vs 7 | 5 vs 8 | 6 vs 9 | 7 vs 10 | 8 vs 1 | 9 vs 2 | 10 vs 3 |
| --- | --- | --- | --- | --- | --- | --- | --- | --- | --- | --- |
| 1 vs 4 |  | t = 2.95  p = 0.01 | t = 3.33  p < 0.01 | t = 4.83  p < 0.001 | t = 5.37  p < 0.001 | t = 1.92  p = 0.09 | t = 2.21  p = 0.06 | t = -3.76  p < 0.01 | t = -2.04  p = 0.07 | t = -3.38  p < 0.01 |
| 2 vs 5 |  |  | t = 0.06  p = 0.96 | t = 2.04  p = 0.07 | t = 1.57  p = 0.16 | t = -0.65  p = 0.60 | t = -0.54  p = 0.63 | t = -6.44  p < 0.001 | t = -4.42  p < 0.001 | t = -4.81  p < 0.001 |
| 3 vs 6 |  |  |  | t = 2.14  p = 0.06 | t = 1.69  p = 0.13 | t = -0.71  p = 0.57 | t = -0.58  p = 0.63 | t = -5.75  p < 0.001 | t = -3.78  p < 0.01 | t = -4.50  p < 0.001 |
| 4 vs 7 |  |  |  |  | t = -0.16  p = 0.89 | t = -2.66  p = 0.02 | t = -2.64  p = 0.02 | t = -6.54  p < 0.001 | t = -4.70  p < 0.001 | t = -5.19  p < 0.001 |
| 5 vs 8 |  |  |  |  |  | t = -2.41  p = 0.04 | t = -1.89  p = 0.09 | t = -6.78  p < 0.001 | t = -5.39  p < 0.001 | t = -6.34  p < 0.001 |
| 6 vs 9 |  |  |  |  |  |  | t = 0.21  p = 0.87 | t = -3.79  p < 0.01 | t = -2.38  p = 0.04 | t = -3.17  p < 0.01 |
| 7 vs 10 |  |  |  |  |  |  |  | t = -4.39  p < 0.001 | t = -2.65  p = 0.02 | t = -3.14  p < 0.001 |
| 8 vs 1 |  |  |  |  |  |  |  |  | t = 1.77  p = 0.11 | t = 1.19  p = 0.30 |
| 9 vs 2 |  |  |  |  |  |  |  |  |  | t = -0.54  p = 0.63 |
| 10 vs 3 |  |  |  |  |  |  |  |  |  |  |

Table S4. Comparisons between queried temporal distance memory pairs in Experiment 2

| Position | 1 vs 4 | 2 vs 5 | 3 vs 6 | 4 vs 7 | 5 vs 8 | 6 vs 9 | 7 vs 10 | 8 vs 1 | 9 vs 2 | 10 vs 3 |
| --- | --- | --- | --- | --- | --- | --- | --- | --- | --- | --- |
| 1 vs 4 |  | t = 7.67  p < 0.001 | t = 5.77  p < 0.001 | t = 4.99  p < 0.001 | t = 5.40  p < 0.001 | t = 7.63  p < 0.001 | t = 8.26  p < 0.001 | t = -8.73  p < 0.001 | t = -8.79  p < 0.001 | t = -8.78  p < 0.001 |
| 2 vs 5 |  |  | t = -4.50  p < 0.001 | t = -3.93  p < 0.001 | t = -4.31  p < 0.001 | t = -0.26  p = 0.87 | t = -0.63  p = 0.65 | t = -9.17  p < 0.001 | t = -8.84  p < 0.001 | t = -8.69  p < 0.001 |
| 3 vs 6 |  |  |  | t = 0.12  p = 0.91 | t = -0.17  p = 0.88 | t = 4.55  p < 0.001 | t = 4.40  p < 0.001 | t = -9.87  p < 0.001 | t = -9.50  p < 0.001 | t = -9.06  p < 0.001 |
| 4 vs 7 |  |  |  |  | t = -0.23  p = 0.88 | t = 4.13  p < 0.001 | t = 3.78  p = 0.001 | t = -9.42  p < 0.001 | t = -9.08  p < 0.001 | t = -8.58  p < 0.001 |
| 5 vs 8 |  |  |  |  |  | t = 4.53  p < 0.001 | t = 4.58  p < 0.001 | t = -9.40  p < 0.001 | t = -8.94  p < 0.001 | t = -8.72  p < 0.001 |
| 6 vs 9 |  |  |  |  |  |  | t = -0.42  p = 0.78 | t = -9.41  p < 0.001 | t = -9.11  p < 0.001 | t = -8.72  p < 0.001 |
| 7 vs 10 |  |  |  |  |  |  |  | t = -9.83  p < 0.001 | t = -9.30  p < 0.001 | t = -9.12  p < 0.001 |
| 8 vs 1 |  |  |  |  |  |  |  |  | t = -0.17  p = 0.88 | t = 0.32  p = 0.84 |
| 9 vs 2 |  |  |  |  |  |  |  |  |  | t = 0.58  p = 0.67 |
| 10 vs 3 |  |  |  |  |  |  |  |  |  |  |
